# The atypical cadherin CELSR2 regulates distinct epithelial–mesenchymal states and oxidative phosphorylation in triple-negative breast cancer

**DOI:** 10.64898/2026.09.15.751697

**Authors:** Julia Koren, Nour-Islem Yanis Saidani, Abdessamad Elkaoutari, Avais Daulat, Pascal Finetti, Emmanuelle Josselin, Stéphane Audebert, Luc Camoin, Roxanne A Gauthier, Rémy Castellano, Arthur Géraud-Crémieux, Fabienne Lamballe, François Bertucci, Flavio Maina, Jean-Paul Borg, Alexandra Walton

**Affiliations:** Aix Marseille Univ, INSERM, CNRS, Institut Paoli-Calmettes, Centre de Recherche en Cancérologie de Marseille (CRCM), Team ‘Targeting signaling networks and microenvironment in cancer’, Marseille, France; Aix Marseille Univ, CNRS, INSERM, Institut Paoli-Calmettes, Centre de Recherche en Cancérologie de Marseille (CRCM), Predictive Oncology laboratory, Label Ligue contre le Cancer, Marseille, France; Aix Marseille Univ, CNRS, INSERM, Institut Paoli-Calmettes, Centre de Recherche en Cancérologie de Marseille (CRCM), TrGET preclinical platform, Marseille, France; Aix Marseille Univ, INSERM, CNRS, Institut Paoli-Calmettes, Centre de Recherche en Cancérologie de Marseille (CRCM), Marseille Proteomics, Marseille, France; Institut National de La Recherche Scientifique, INRS-Centre Armand-Frappier Santé Biotechnologie, Université du Québec, Laval, QC, Canada; Université Clermont Auvergne, Faculté de Médecine, F-63000 Clermont-Ferrand France; Aix Marseille Univ, CNRS, INSERM, Institut Paoli-Calmettes, Centre de Recherche en Cancérologie de Marseille (CRCM), Department of Medical Oncology, Marseille, France; Aix Marseille Univ, CNRS, INSERM, Institut Paoli-Calmettes, Centre de Recherche en Cancérologie de Marseille (CRCM), COMPO Team, Marseille, France; Aix Marseille Univ, Turing Center for Living Systems, Marseille, France; Institut Universitaire de France, Paris, France

**Keywords:** Adhesion G protein-coupled receptor, CELSR2, triple-negative breast cancer, epithelial–mesenchymal plasticity, intratumoral heterogeneity, oxidative phosphorylation

## Abstract

Triple-negative breast cancer (TNBC) is characterized by marked intratumoral heterogeneity, which contributes to metastatic dissemination, therapeutic resistance, and poor clinical outcome. Epithelial-mesenchymal plasticity is a major contributor to this heterogeneity. Identifying the molecular regulators governing epithelial-mesenchymal plasticity is therefore essential to understanding how distinct tumor cell populations arise and fuel TNBC progression.

Here, we identify the adhesion G protein-coupled receptor CELSR2 as an independent poor-prognosis marker and regulator of epithelial-mesenchymal plasticity in TNBC. Spatial transcriptomic analyses revealed segregation of CELSR2^high^ tumor cells within the epithelial tumor core and CELSR2^low^ tumor cells at the invasive front. In line with the latter observation, CELSR2 expression decreases during EMT and its loss promotes acquisition of a hybrid epithelial-mesenchymal state. CELSR2^high^ tumor cells display a CREB-associated oxidative phosphorylation (OXPHOS) program and promote cell proliferation and primary tumor growth. Conversely, CELSR2^low^ cells display features of a more migratory phenotype.

Collectively, our findings identify CELSR2 as a previously unknown regulator of epithelial-mesenchymal plasticity and reveal that heterogeneous CELSR2 expression shapes spatially and functionally distinct tumor cell populations, thereby contributing to intratumoral heterogeneity in TNBC.

**Statement of Significance:** Heterogeneous CELSR2 expression contributes to the coexistence of spatially and functionally distinct tumor cell populations characterized by different epithelial-mesenchymal states and oxidative metabolic properties, thereby promoting intratumoral heterogeneity in TNBC.

## Introduction

Triple-negative breast cancer (TNBC) is the most aggressive subtype of breast cancer and remains associated with poor clinical outcome due to its high propensity for recurrence, metastatic dissemination, and therapeutic resistance^1^. A major challenge in TNBC is its pronounced intratumoral heterogeneity, which complicates therapeutic management and contributes to disease progression^2–4^. Among the processes contributing to this heterogeneity, epithelial-mesenchymal plasticity, the reversible ability of tumor cells to dynamically switch between epithelial, hybrid epithelial-mesenchymal, and mesenchymal states, has emerged as a major source of phenotypic diversity^5^. Epithelial-to-mesenchymal transition (EMT), an evolutionarily conserved developmental program frequently reactivated by cancer cells, promotes the acquisition of migratory and invasive properties^6^. Rather than a binary switch, EMT is now viewed as a continuum of epithelial, hybrid, and mesenchymal states that coexist within the same tumor^7^. Notably, hybrid states have been associated with enhanced metastatic potential and increased cellular plasticity^7^. Reacquisition of epithelial features through mesenchymal-to-epithelial transition (MET) is required for metastatic outgrowth at distant sites^8,9^. Together, these observations highlight that the ability of tumor cells to reversibly transition between epithelial and mesenchymal states is a critical determinant of tumor progression. Identifying the molecular regulators governing epithelial-mesenchymal plasticity is therefore essential to understanding how distinct cellular populations arise and contribute to TNBC progression.

Among the signaling pathways implicated in TNBC, increasing evidence supports a role for non-canonical WNT/Planar Cell Polarity (PCP) signaling in several processes associated with EMT, including tumor invasion, metastasis, and treatment resistance^10–12^. The WNT/PCP pathway relies on a set of conserved membrane proteins that coordinate cell polarity, cell-cell interactions, and tissue organization in physiological and disease contexts^13^. Among these proteins, Cadherin EGF LAG seven-pass G-type receptor 2 (CELSR2) is a large atypical cadherin and seven-pass transmembrane receptor belonging to the adhesion G protein-coupled receptor (aGPCR) family. CELSR2 was initially characterized for its role in neurodevelopment, where it contributes to neuronal migration, axon guidance and synaptic connectivity^14,15^. More recently, dysregulated CELSR2 expression has been linked to cancer progression in hepatocellular carcinoma and glioma, and proposed as a prognostic biomarker and a potential therapeutic target^16,17^. In addition, increased CELSR2 expression has been reported in experimental models of mammary tumorigenesis^18^. However, whether CELSR2 is associated with specific tumor-cell states or functional programs in breast cancer, particularly in TNBC, remains unknown.

Here, we identify CELSR2 as an important factor in TNBC intratumoral heterogeneity and demonstrate that variations of CELSR2 expression define distinct epithelial-mesenchymal programs. CELSR2^high^ cells maintain epithelial characteristics in the tumor core and a CREB-associated oxidative phosphorylation program related to proliferation and tumor growth. In contrast, loss of CELSR2 at the invasive front correlates with decreased cell-matrix adhesion and emergence of a hybrid EMT state, marked by preserved epithelial cell-cell junctions and enhanced migratory properties. CELSR2 expression is reduced in circulating tumor cells and re-expressed in metastatic lesions. *In vivo* experiments confirm a key role of CELSR2 in tumor growth. Together, our findings identify CELSR2 as a previously unrecognized regulator of epithelial-mesenchymal plasticity in TNBC and highlight its role in intratumoral heterogeneity and tumor progression.

## Materials and Methods

### Cell lines and cell culture

Human breast cancer cell lines (MDA-MB-231, MDA-MB-468, and MCF-7) and HEK293T cells were obtained from the American Type Culture Collection (ATCC; Manassas, VA, USA). Cells were cultured in high-glucose Dulbecco’s modified Eagle’s medium (DMEM; #41965, Thermo Fisher Scientific) supplemented with 10% heat inactivated fetal bovine serum (FBS), 100 U/mL penicillin, and 100 μg/mL streptomycin (#15140-122, Thermo Fisher Scientific) and maintained at 37 °C in a humidified atmosphere containing 5% CO₂. Mouse Mammary Gland Tumor (MGT) cell lines were derived from independent mammary tumors arising in the MMTV-R26^Met^ mouse model using the procedure reported by Lamballe et al ^19^. Cells were cultured in the complete medium described by Lamballe et al. and maintained at 37 °C in a humidified atmosphere containing 5% CO₂. All cell lines were routinely tested and confirmed to be free of mycoplasma contamination.

### siRNA-mediated knockdown

Cells were reverse-transfected with 20 nM of individual siRNAs using Lipofectamine™ RNAiMAX (#13778075, Thermo Fisher Scientific). Cells were harvested 48 h later for RNA extraction. A non-targeting siRNA (Dharmacon) was used as a negative control. The sequences of the siRNAs used in this study are provided in **Supplementary Table 1**.

### Plasmid transfection

Cells were transfected with 1.5 μg of plasmid DNA using Lipofectamine™ LTX and Plus Reagent (#15338100, Thermo Fisher Scientific) according to the manufacturer’s instructions. Plasmids used in this study are listed in **Supplementary Table 2**.

### Lentiviral production and transduction

Lentiviral vectors encoding mCherry, mCh-T2A-CELSR2, shCELSR2 and shCtrl (**Supplementary Table 2 and Table 3**) were produced in HEK293T cells by co-transfection of the lentiviral transfer plasmid with the packaging plasmid psPAX2 and the envelope plasmid pMD2.G (VSV-G) using Lipofectamine™ LTX and Plus Reagent (#15338100, Thermo Fisher Scientific). Viral supernatants were collected 72 h after transfection, filtered through 0.22-μm membranes, and concentrated using Lenti-X Concentrator (#631231, Takara). Target cells were transduced for 72 h. For shRNA-mediated knockdown, transduction was performed in the presence of polybrene, whereas protamine sulfate (10 μg/mL) was used for mCherry and mCh-T2A-CELSR2 vectors. Stable cell populations expressing mCherry or mCh-T2A-CELSR2 were isolated by fluorescence-activated cell sorting (FACS), whereas stable shRNA-expressing cells were selected with puromycin (1 μg/mL) for 2 weeks.

### Quantitative reverse transcription-PCR

Total RNA was extracted using the RNeasy Mini Kit (#74104, Qiagen) according to the manufacturer’s instructions. cDNA was synthesized from 1 μg of mRNA using the iScript ReverseTranscription Supermix (#L001404, Bio-Rad, USA). Quantitative PCR was performed using SYBR Green Master Mix (#56475, Thermo Fisher Scientific, Carlsbad, USA) on a CFX96 Real-Time PCR Detection System (Bio-Rad) using CFX manager software. Relative gene expression was calculated using the ΔΔCt method, with GAPDH as the internal control. Primer sequences are listed in **Supplementary Table 4**.

### Cell proliferation assay

The Cell Counting Kit-8 assay (CCK-8, #C0005, TargetMol, Boston, USA) was used to indirectly assess cell proliferation. Cells were seeded in 96 well-plates at 1 × 10³ cells/well (MGT19, MGT20, and MDA-MB-231) or 2 × 10³ cells/well (MDA-MB-468). After 72h, CCK-8 reagent (10% v/v) was added to each well and incubated for 2 h at 37 °C. Absorbance was measured at 450 nm using a microplate reader after subtraction of the background signal.

### Crystal violet assay

Forty-eight hours after seeding, cells were fixed with 4% paraformaldehyde (#28908, Thermo Fisher Scientific) and stained with 1% crystal violet. After washing, the bound dye was solubilized in 1% sodium dodecyl sulfate (SDS), and absorbance was measured at 570 nm using a microplate reader after background subtraction.

### Western Blot analysis

Cells were lysed in ice-cold lysis buffer (50 mM HEPES-NaOH, pH 8.0, 150 mM NaCl, 10% glycerol, 2 mM EDTA, and 0.5% NP-40) supplemented with protease and phosphatase inhibitor cocktails (Sigma-Aldrich). Protein concentrations were determined using the Pierce BCA Protein Assay Kit (#23225, Thermo Fisher Scientific). Equal amounts of protein were loaded on a NuPAGE 4–12% Bis-Tris gel (#NP0336BOX, Invitrogen) and transferred onto 0.45-μm nitrocellulose membranes (#10600002, Cytiva). Membranes were stained with Ponceau S, blocked in TBS containing 0.1% Tween-20 and 5% non-fat dry milk, and incubated overnight at 4 °C with primary antibodies. After washing, membranes were incubated with HRP-conjugated secondary antibodies (Thermo Fisher Scientific) for 1 h at room temperature. HRP mediated chemiluminescence was detected using ECL Prime or ECL Select Western Blotting Detection Reagents (#RPN2235 or #RPN2236, Cytiva) and visualized with a G:BOX imaging system (Syngene). Band intensities were quantified using ImageJ. Antibodies and dilution factors are listed in **Supplementary Table 5**.

### Immunofluorescence staining and confocal microscopy

Cells grown on collagen I-coated glass coverslips were fixed with 4% paraformaldehyde (#28908, Thermo Fisher Scientific), permeabilized with 0.4% Triton X-100 in PBS, and blocked with 3% BSA in PBS. Cells were incubated overnight at 4 °C with the indicated primary antibodies, washed, and incubated with the appropriate Alexa Fluor-conjugated secondary antibodies. Coverslips were mounted using ProLong™ Gold Antifade Mountant with DAPI (#P36931, Invitrogen). Images were acquired using a Zeiss LSM880 confocal laser scanning microscope. Antibodies and dilution factors are listed in **Supplementary Table 5**.

### CREB luciferase reporter assay

HEK293T cells were co-transfected with GFP or GFP-CELSR2 expression plasmids together with a CRE-responsive Firefly luciferase reporter plasmid and a Renilla luciferase control plasmid (**Supplementary Table 2**) using Lipofectamine™ LTX and Plus Reagent (#15338100, Thermo Fisher Scientific) according to the manufacturer’s instructions. 48 hours after transfection, cells were lysed in Passive Lysis Buffer 1X (#E194A, Promega). Firefly and Renilla relative luminescence units (RLU) were determined using the Dual-Glo Luciferase Reporter Assay System (#E1910, Promega) according to the manufacturer’s instructions. Firefly luciferase activity was normalized to Renilla luciferase activity, and the resulting Firefly/Renilla ratios were expressed as fold induction relative to GFP-transfected cells.

### Cell-matrix adhesion assay

Six-well plates were coated overnight at 4 °C with collagen I (#66914600, Roche) or fibronectin (10 μg/mL each) and blocked with 1% BSA in PBS for 1 h at room temperature. Cells (9 x 10^4^ per well) were seeded in serum-free medium. At the indicated time points (15, 30, 60, and 120 min), non-adherent cells were removed, and adherent cells were fixed with 4% formaldehyde, stained with 1% crystal violet, and the bound dye was solubilized in 1% SDS. Absorbance was measured at 570 nm using a microplate reader after background subtraction.

### Single-cell tracking

MDA-MB-231 cells (5 x 10^4^) were seeded on collagen I-coated 6-well plates. The following day, live-cell imaging was performed using an Olympus CellVivo IX83 microscope controlled with cellSens software. Images were acquired every 10 min for 24 h. Cell trajectories were manually tracked using the Manual Tracking plugin in Fiji/ImageJ. Migration speed was calculated as the total distance traveled divided by the duration of migration, whereas Euclidean distance was defined as the linear distance between the initial and final cell positions.

### Mitochondrial oxygen consumption rate (OCR)

Oxygen consumption rate (OCR) was measured using a Seahorse XFe24 Extracellular Flux Analyzer (#00420148, Agilent Technologies, Santa Clara, CA, USA). MDA-MB-468 cells transfected with si*CELSR2* or non-targeting siRNA were seeded (4 x 10^4^ cells/well) in Seahorse XFe24 microplates (#102340-100, Agilent Technologies). Before analysis, cells were incubated in Seahorse XF DMEM (#103575-100, Agilent Technologies) supplemented with 2 mM glutamine, 1 mM sodium pyruvate, and 10 mM glucose according to the manufacturer’s instructions. OCR was measured following sequential injections of oligomycin (1 μM), FCCP (1 μM), and rotenone/antimycin A (0.5 μM each). At the end of the assay, cells were fixed and stained with crystal violet for cell number normalization. Basal respiration, maximal respiration, spare respiratory capacity, and non-mitochondrial respiration were calculated using Wave software (Agilent Technologies).

### Animal studies

All animal experiments were performed in agreement with the French Guidelines for animal handling and approved by the local ethic committee (C2EA14) for Animal Experimentation (Agreement no. APAFIS#33446--2021092811105354). NOD/SCID/γc null mice (NSG) were obtained from Dr. C. Rivers (Margate, UK). Mice were housed under sterile conditions with sterilized food and water provided *ad libitum* and maintained on a 12-h light and 12-h dark cycle. MDA-MB-468 (1 × 10^6^cells/mouse) in a 1/2 (v/v) Matrigel (Becton Dickinson Bioscience) suspension were injected into mammary fat pads of 9-10-week-old female NSG mice. Tumor growth was monitored by digital caliper measurements and by calculating volumes (length × width × height × pi/6). To compare tumor size between the experimental groups, two-way ANOVA followed by Tukey’s multiple-comparisons test was used.

### LC/MS

Total proteomes from CELSR2-depleted cells (MDA-MB468 si*CELSR2* and MCF7 si*CELSR2*) were compared with their respective control cells (MDA-MB468 siCtrl and MCF7 siCtrl) by label-free quantitative mass spectrometry analysis. 15µg of cell lysates were loaded on NuPAGE™ 4–12% Bis–tris acrylamide gels according to the manufacturer’s instructions (Invitrogen, Life Technologies). Running of samples was stopped as soon as proteins stacked as a single band and following imperial blue staining (Life Technologies), the upper part of the gel containing the proteins was cut and processed for classical in gel digestion (washes, thiols reduction with 10 mM DTT and cystein alkylation with 55 mM iodoacetamide). Each band was further digested as previously described with trypsin and analyzed by liquid chromatography (LC)-tandem MS (MS/MS) using an Orbitrap Fusion Lumos Tribrid Mass Spectrometer (ThermoFisher Scientific, San Jose, CA) online with a nanoRSLC Ultimate 3000 chromatography system (ThermoFisher Scientific, Sunnyvale, CA). First, peptides were concentrated and purified on a pre-column PepMap100 C18, 2 cm × 100 μm I.D, 100 Å pore size, 5 μm particle size in solvent A (0.1% formic acid in 2% acetonitrile). In the second step, peptides were separated on a reverse phase LC EASY-Spray C18 column PepMap RSLC C18, 50 cm × 75 μm I.D, 100 Å pore size, 2 μm particle size (ThermoFisher) at 300 nL/min flow rate and 40 °C. After column equilibration using 4% of solvent B (20% water - 80% acetonitrile - 0.1% formic acid), peptides were eluted from the analytical column by a two-step linear gradient (2–20% acetonitrile/H₂O; 0.1% formic acid for 90 min and 20–45% acetonitrile/H₂O; 0.1% formic acid for 20 min). For peptide ionization in the EASY-Spray nanosource in front of mass spectrometer, spray voltage was set at 2.2 kV and the capillary temperature at 275 °C. The Orbitrap Lumos was used in data-independent mode with the following parameters. First, MS spectra were acquired in the Orbitrap in the range of m/z 399–1500 at a FWHM resolution of 120,000 measured at 400 m/z. AGC target was set at standard parameters with an automatic Maximum Injection Time. MS2 spectra were acquired in the Orbitrap with a resolution of 30,000, in the mass range of 200–1800 m/z after isolation of parent ion in the quadrupole and fragmentation in the HCD cell under collision Energy of 30%. DIA parent ion range was from 400 to 1000 m/z divided into 40 windows 16 Da wide and from 1000 to 1500 m/z divided into 10 windows 50 Da wide.

### Data Processing Protocol

Relative intensity-based label-free quantification (LFQ) was processed using the DIA-NN 1.8 algorithm. Raw files were searched against the Human database extracted from UniProt on the 10th of January 2023 and containing 20404 entries (reviewed) with the addition of a protein contaminant bank^20^. The following parameters were used for searches: (i) trypsin allowing cleavage before proline; (ii) one missed cleavage was allowed; (iii) cysteine carbamidomethylation (+57.02146) as a fixed modification and methionine oxidation (+15.99491) and N-terminal acetylation (+42.0106) as variable modifications; (iv) a maximum of 1 variable modification per peptide allowed; and (v) minimum peptide length was 7 amino acids and a maximum of 30 amino acids. The match between runs option was enabled to transfer identifications across different LC-MS/MS replicates based on their masses and retention time. The precursor false discovery was set to 1%. DIA-NN parameters were set on Single-pass mode for Neural Network classifier, Robust LC High precision for quantification strategy and RT-dependent mode for Cross-run normalization. Library was generated using Smart profiling set up. The main output file from DIA-NN was further filtered at 1% FDR and LFQ intensity was calculated using our DIAgui package at 1% q-value (https://github.com/marseille-proteomique/DIAgui)^21^.

The statistical analysis was done with Perseus program (version 1.6.15.0) from the MaxQuant environment (www.maxquant.org)^22^. Quantifiable proteins were defined as those detected in above 70% of samples in one condition or more. Missing values were replaced using data imputation by randomly selecting from a normal distribution centered on the lower edge of the intensity values that simulates signals of low abundant proteins using default parameters (a downshift of 1.8 standard deviation and a width of 0.3 of the original distribution). To determine whether a given detected protein was specifically differential, a two-sample t-test was done using permutation-based FDR-controlled at 0.01 and employing 250 permutations. The p value was adjusted using a scaling factor s0 with a value of 1. Analysis was done on biological triplicates, each injected twice on mass spectrometers.

### Bulk transcriptomic datasets

We analyzed our breast cancer gene expression database^23^ pooled from 36 public data sets (**Supplementary Table 6**), comprising 8,982 non-redundant non-metastatic, non-inflammatory, primary, and invasive breast cancer samples, and 429 tumor-adjacent normal breast tissues (AT). For each sample, both the gene expression profile generated using DNA microarrays or RNA-Seq and the clinicopathological annotations were available. These sets had been gathered from the National Center for Biotechnology Information (NCBI)/Genbank GEO, ArrayExpress, European Genome-Phenome Archive, The Cancer Genome Atlas portal (TCGA) databases, and authors’ website. We also included the gene expression data of 179 normal breast tissues from healthy people, obtained from the UCSC Xena Browser (GTEx). The pre-analytic data processing was done as previously reported^23^. We collected the DNA copy number variation (CNV) data of breast cancer samples from TCGA and expressed as log₂(tumor/normal). *CELSR2* expression in tumors was analysed as continuous value; associations between *CELSR2* mRNA expression and CNV were assessed using Pearson correlation. Correlations between *CELSR2* expression and expression of “mesenchymal” or “epithelial” genes of interest were evaluated using the Spearman correlation analysis. We also applied to each dataset separately the Taube et al’s signature^24^. This signature is a transcriptional gene-expression program derived from the common gene-expression changes induced by multiple independent epithelial-to-mesenchymal transition (EMT) stimuli in human mammary epithelial cells: it contains 246 genes, including 87 up-regulated, and 159 down-regulated during EMT. This signature is used to quantify EMT phenotype in breast cancer. For each cancer sample, we computed an EMT metagene score from this signature as the difference between the mean expression of up-regulated and down-regulated genes, after median-centering of each gene within each study separately. This score was analysed as continuous value. Linear regression (glm) was used to evaluate association between *CELSR2* expression and EMT score, in all TNBC samples and within each dataset, yielding an Geometric Mean Ratio (GMR) with its 95 % confidence interval as the effect size for each dataset. A meta-analytic approach was conducted using the R package meta where the common-effect model was applied to pool the 14 dataset-specific GMRs. Heterogeneity among datasets was quantified using Cochran’s Q test, the I² statistic, and the between-study variance τ² (DerSimonian–Laird estimator). Our breast cancer cohort of 8,982 patients included 3,454 informative for MFS. Samples were stratified into low and high *CELSR2* expression groups by fitting a 2-component Gaussian mixture model (GMM) *via* maximum likelihood estimation as previously reported^25^ Kaplan-Meier curves were compared using the two-sided log-rank tests, and hazard ratios were estimated using Cox proportional hazards models. Uni- and multivariate analyses for MFS were performed using Cox regression analysis (Wald test). The variables submitted to univariate analyses included patients’ age at diagnosis, pathological tumor type, grade and size, athological axillary node status (pN), delivery of adjuvant chemotherapy, and CELSR2 expression group. Multivariate analyses included the variables with a *P*-value inferior to 0.10 in univariate analysis. Two other publicly available cohorts were used : GSE209998 to compare primary and metastatic BC tumors, and GSE111842 to compare CELSR2 expression between circulating tumor cells (CTCs) and primary BC tumors. Differences were evaluated using two-sided Wilcoxon rank-sum tests.

### Single-cell RNA sequencing analysis

Single-cell RNA-seq data from TNBC tumors (GSE161529; 54,820 cells from eight patients) were processed using Seurat. Quality control filtering was performed based on the number of detected genes per cell and the proportion of mitochondrial transcripts. Data were normalized, highly variable genes were identified, and expression values were scaled prior to dimensionality reduction by principal component analysis (PCA) and visualization using t-distributed stochastic neighbor embedding (t-SNE). *CELSR2* and *ZEB1* expression were visualized using log-normalized values. Cells were considered positive for a given gene when normalized expression was greater than zero. Epithelial and mesenchymal module scores were computed using curated gene signatures and Seurat’s module scoring *AddModuleScore()* function. Differences between CELSR2-positive and -negative tumor cells were evaluated using Wilcoxon rank-sum tests.

### Spatial transcriptomics analysis

Spatial transcriptomic data were downloaded from Wang et *al* study^26^ and analyzed using the BCTL-Bordet/ST pipeline provided by the authors (https://github.com/BCTL-Bordet/ST/). Processed expression matrices and spatial coordinates were used for downstream analyses. Spots were annotated according to the histopathological classifications provided in the original study, and analyses were restricted to tumor-associated spots. To distinguish tumor core from invasive front regions, a k-nearest neighbor graph (k = 6) was constructed using spatial coordinates. For each tumor spot, the proportion of neighboring non-tumor spots was calculated. Spots with at least 25% non-tumor neighbors were classified as peripheral (invasive front), while others were classified as central. Binary detection of CELSR2 expression was defined as normalized expression greater than zero. For each patient, the proportion of CELSR2-positive tumor spots was calculated separately in central and peripheral regions. Differences were evaluated using *Fisher*’s exact tests, and odds ratios with 95% confidence intervals were calculated to estimate enrichment or depletion at the invasive front. Patient-level paired comparisons were assessed using Wilcoxon signed-rank tests. To assess spatial co-expression between CELSR2 and EMT-associated genes, spot-level binary detection was performed for each gene using a threshold of normalized expression greater than zero. For each EMT marker, spots were classified into four categories: CELSR2-only positive, EMT-only positive, double-positive, or double-negative.

### ChIP-seq analysis

Publicly available ChIP-seq datasets for ZEB1 in MDA-MB-231 cells (GEO GSE89203) and pCREB in MDA-MB-134 cells (GEO GSE109103), together with TWIST1 ChIP-exo data generated in MCF-7 cells (GEO GSE189826), were downloaded from the Gene Expression Omnibus (GEO). Processed normalized signal tracks (bigWig files) and, when available, corresponding peak calls provided by the original studies were used for analysis. Genomic occupancy profiles were visualized using the Integrative Genomics Viewer (IGV; Broad Institute) at the CELSR2, VIM, CDH1, ATP5A1, ATP5H, NDUFV2, and COX1 loci. Gene models, exon structures, and annotated translation start sites (ATG) were displayed according to the reference genome assembly used in the original studies. All genomic visualizations were generated from the processed signal tracks without additional read alignment, normalization, or peak calling.

### Statistical analysis

Statistical analyses were performed using R or GraphPad Prism version 8.0 (GraphPad Software). Appropriate statistical tests were applied as indicated in the corresponding figure legends. Data are presented as mean ± SD or SEM, as indicated. *P* < 0.05 was considered statistically significant. When applicable, *P* values were adjusted for multiple testing using the Benjamini–Hochberg method.

## Data and code availability

The mass spectrometry proteomics data have been deposited in the ProteomeXchange Consortium via the PRIDE partner repository under the dataset accession numbers PXD082952 and PXD082953, corresponding to the MCF-7 and MDA-MB-468 datasets, respectively.

Reviewer access:

Reviewers can access the datasets through the PRIDE website using the following project accessions and reviewer tokens:

- MCF-7: Project accession PXD082952
- MDA-MB-468: Project accession PXD082953

## Results

### High *CELSR2* expression in TNBC is associated with poor prognosis and epithelial features

To investigate the clinical relevance of CELSR2 in breast cancer, we first analyzed its expression in our large base of publicly available transcriptomic datasets. *CELSR2* expression was significantly increased in breast tumors compared with healthy breast tissues and cancer-adjacent tissues (AT) (**Fig. 1A**). Furthermore, *CELSR2* mRNA expression positively correlated with copy-number gain in breast cancer samples, suggesting that genomic amplification may contribute in part to its transcriptional upregulation (**Fig. 1B**).

**Fig. 1.**
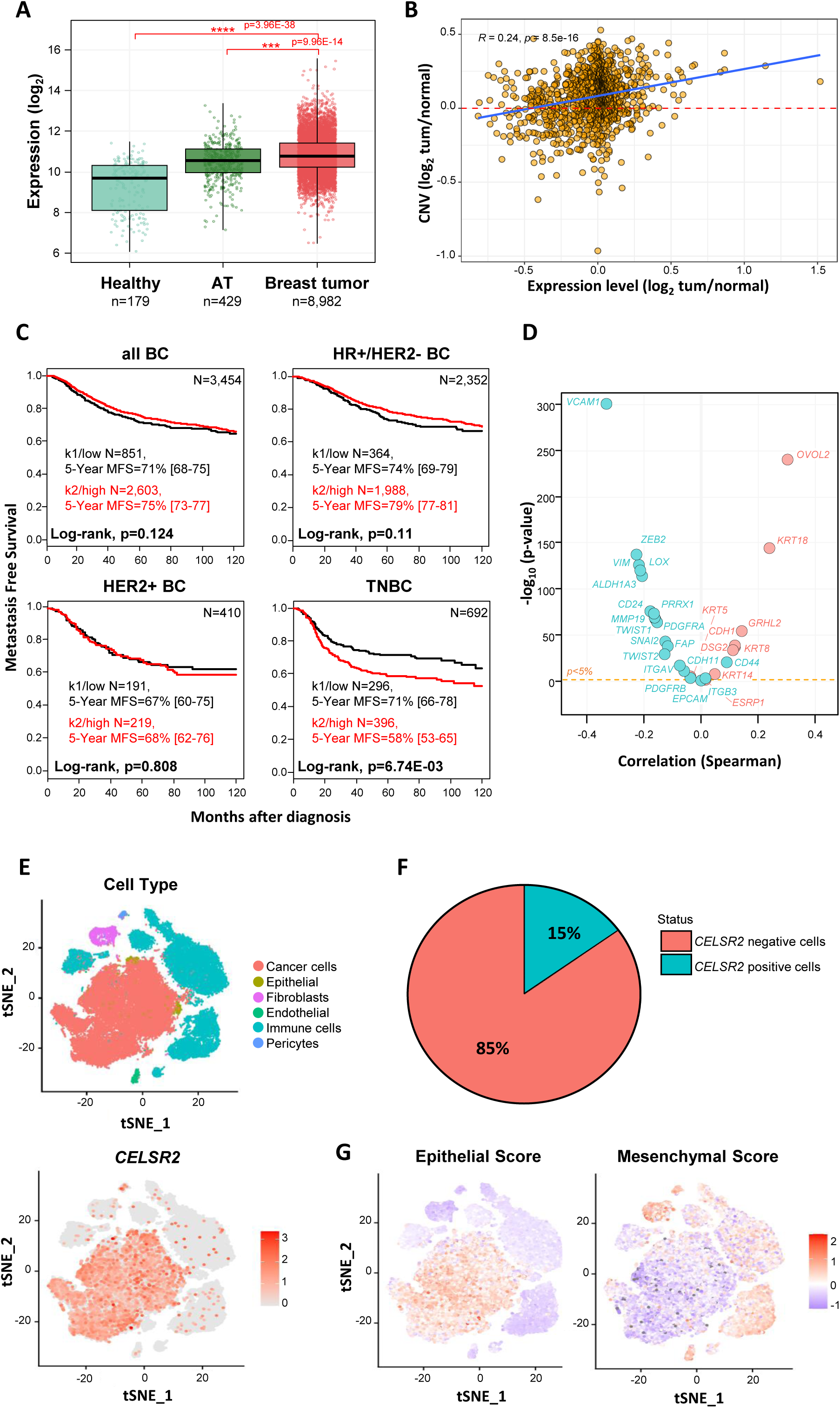
High *CELSR2* expression in TNBC is associated with poor prognosis and epithelial features. (**A**) *CELSR2* mRNA expression in healthy breast tissues (GTEx), cancer-adjacent normal breast tissues (AT), and primary breast tumors from our gene expression database (n = 179, 429, and 8,982 respectively). Expression values are shown as normalized log₂-transformed values. Box plots indicate the median and interquartile range; statistical significance was determined using the two-sided Student *t*-test. (**B**) Correlation between *CELSR2* mRNA expression and copy number variation (CNV) in breast cancer samples (n = 1,104). CNV values are expressed as log₂(tumor/normal). Each dot represents one tumor sample, and the blue line indicates the linear regression fit. Pearson correlation analysis demonstrates a positive association between *CELSR2* expression and gene copy number (CNV). (**C**) Kaplan–Meier curves of metastasis-free survival (MFS) according to *CELSR2* expression in the overall breast cancer cohort (BC) and in molecular subtypes (HR+/HER2−, HER2+, and TNBC) using a combined cohort composed of Institut Paoli-Calmettes (IPC) patients and publicly available breast cancer datasets. Patients were stratified into low- and high-*CELSR2* expression groups. Statistical significance was assessed using the log-rank test. (**D**) Correlation of *CELSR2* expression with expression of epithelial- and mesenchymal-associated genes in TNBC samples of our gene expression database. Each dot represents one gene, plotted according to its Spearman correlation coefficient with *CELSR2* expression (x-axis) and statistical significance (−log₁₀ *P* value; y-axis). The dashed line indicates the significance threshold. Representative epithelial and mesenchymal markers are annotated, highlighting the association of CELSR2 with epithelial transcriptional programs. (**E**, **G**) t-distributed stochastic neighbor embedding (t-SNE) visualization of single-cell RNA sequencing data from TNBC tumors (GSE161529; 54,820 cells from 8 patients). (**E**) Cell-type annotation (top) and *CELSR2* expression (bottom). (**F**) Proportion of *CELSR2*-positive and *CELSR2*-negative tumor cells identified in the TNBC single-cell dataset (GSE161529). The pie chart indicates the percentage of tumor cells expressing *CELSR2*. (**G**) Projection of epithelial and mesenchymal scores onto the same embedding.

We next evaluated the prognostic significance of *CELSR2* expression in the 3,454 patients from our cohort informative for MFS. While high *CELSR2* levels were not significantly associated with MFS in the overall breast cancer population, HER2-positive (HER2+) patients, or hormone receptor-positive/HER2-negative (HR+/HER2−) patients, high *CELSR2* expression was significantly associated with shorter MFS specifically in TNBC patients (**Fig. 1C**), indicating a subtype-specific prognostic value in TNBC. The 5-year MFS was 71% (95%CI, 66-78) in the CELSR2-low group and 58% (CI95% 53-65) in the CELSR2-high group (*P* = 6.74E-03, log-rank test). Uni- and multivariate prognostic analyses are shown in **Supplementary Table 7**. In multivariate analysis, CESLR2-high group remained an independent poor-prognosis variable (*P* = 2.65E-02, Wald test).

To further characterize the transcriptional signatures associated with *CELSR2* expression in this TNBC cohort of our gene expression database, we analyzed the correlation between its expression and those of established epithelial or mesenchymal markers. We found that *CELSR2* expression negatively correlated with mesenchymal markers and positively with epithelial markers (**Fig. 1D**), supporting an association between *CELSR2* expression and epithelial features in TNBC tumors. Single-cell RNA sequencing analyses of a publicly available TNBC cohort (GSE161529) showed that *CELSR2* expression is predominantly detected in tumor cells rather than in stromal cell populations (**Fig. 1E**). Among tumor cells, approximately 15% were classified as CELSR2-positive (**Fig. 1F**). Notably, *CELSR2*-positive tumor cells displayed higher epithelial scores and lower mesenchymal scores compared with *CELSR2*-negative cells (**Fig. 1G**), strengthening the association between *CELSR2* expression and an epithelial transcriptional signature in TNBC.

We then took advantage of a panel of murine TNBC mammary gland tumor (MGT) cell lines, derived from individual tumors from the previously described *MMTV-R26^Met^* TNBC model^19^. CELSR2 protein levels varied according to the epithelial or mesenchymal phenotype of the cell lines, as defined by the expression of E-Cadherin and the mesenchymal markers N-cadherin and Vimentin (**Fig. S1A**). Epithelial-like MGT cell lines expressed detectable levels of CELSR2 in contrast to mesenchymal-like cell lines (**Fig. S1A**). Because the antibody used for western blot analyses recognizes the N-terminal region of CELSR2, which has previously been reported to undergo proteolytic cleavage^27^, RT-qPCR analyses were also performed with 2 sets of primers, and confirmed reduced *CELSR2* mRNA levels in mesenchymal cell lines (**Fig. S1B**).

Together, these findings identify CELSR2 as a clinically relevant prognostic marker in TNBC. High *CELSR2* levels are associated with epithelial features and poor prognosis. Thus, CELSR2^high^ epithelial tumor cells may contribute to TNBC progression alongside mesenchymal cell populations traditionally associated with aggressive disease.

### CELSR2 expression decreases during EMT and regulates epithelial-mesenchymal plasticity in TNBC cells

Based on our observations linking low CELSR2 expression with mesenchymal features in TNBC, we investigated whether CELSR2 is associated with the EMT program. We analyzed the correlation between *CELSR2* expression and the EMT metagene score, a transcriptomic score reflecting mesenchymal features. We observed a significant negative association in our series of 747 TNBC samples (OR=0.88, p=3.18E-03, **Fig. 2A**). Of note, this negative association showed homogeneity across each of the 14 data sets separately, supporting its robustness (common OR=0.86, p= 1.51E-04; **Fig. 2A**, **Fig. S2**). We next examined whether EMT induction modulates CELSR2 expression using MDA-MB-468 TNBC cells repeatedly stimulated with human epidermal growth factor (hEGF) over a 5-day period (**Fig. 2B**). Following hEGF treatment, cells progressively acquired a spindle-like morphology, associated with increased cell dispersion and loss of cell-cell contacts^28^. RT-qPCR analyses showed increased expression of mesenchymal markers while epithelial markers remained expressed, a pattern compatible with a hybrid EMT phenotype (**Fig. 2C**) characterized by the coexistence of epithelial and mesenchymal markers^7^. Western blot analyses further supported this conclusion as marked Vimentin upregulation with partial reduction, but not complete loss, of E-Cadherin expression were observed (**Fig. 2D**). Under these conditions, CELSR2 protein (**Fig. 2D**) and mRNA (**Fig. 2E**) expression levels were significantly reduced upon EMT induction. To further validate these observations, murine epithelial-like MGT14 and MGT20 TNBC cells were treated with TGF-β on days 1 and 3 to induce EMT over a 5-day period (**Fig. S3A**). CELSR2 expression was also significantly reduced following induction of a hybrid EMT state (**Fig. S3B–I**), further supporting the association between hybrid EMT and decreased CELSR2 expression in both human and murine TNBC cells.

**Fig. 2.**
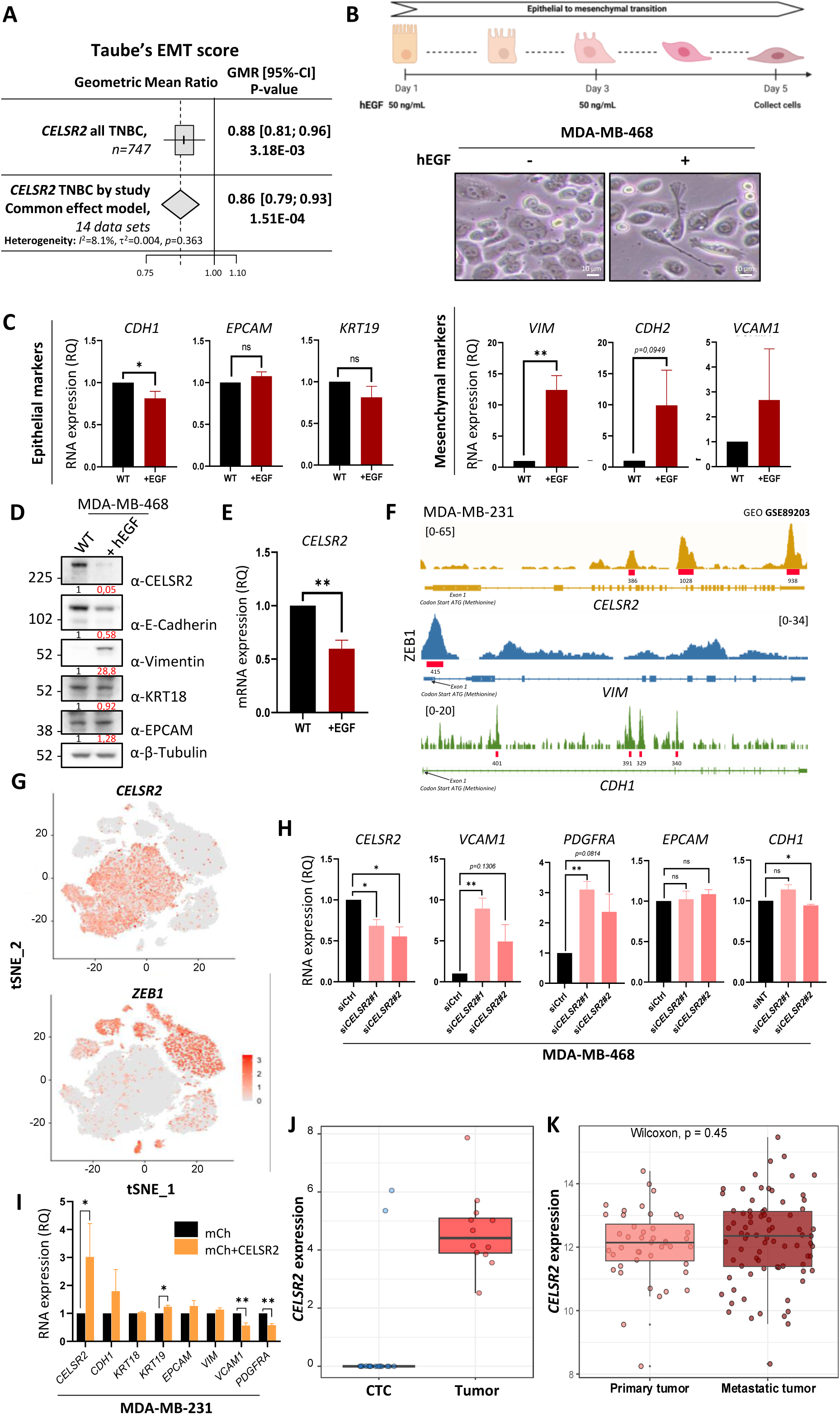
CELSR2 expression decreases during EMT and regulates epithelial-mesenchymal plasticity in breast cancer cells. (**A**) Forest plot showing the negative association between the Taube et al’s EMT metagene score and *CELSR2* mRNA expression in TNBC tumors (n = 747). The forest plots display the Geometric Mean Ratio (GMR) in the whole series (top; n = 747) and the pooled GMR (diamond) at the bottom. Heterogeneity among datasets was quantified using Cochran’s Q test. (**B**) Top: Experimental design for EMT induction in MDA-MB-468 cells. Cells were left untreated (WT) or stimulated with human epidermal growth factor (hEGF, 50 ng/mL) on day 1 and again 72 h later. Cells were harvested on day 5. Bottom: Representative images of untreated (WT) and hEGF-treated MDA-MB-468 cells. (**C**) RT-qPCR analysis of epithelial (left) and mesenchymal (right) marker expression in untreated (WT) and hEGF-treated MDA-MB-468 cells. Gene expression was normalized to *GAPDH* and is presented as relative quantification (RQ). (**D**) Western blot analysis of CELSR2, E-cadherin, KRT18, EPCAM, Vimentin, and β-tubulin in untreated (WT) and hEGF-treated MDA-MB-468 cells. β-Tubulin served as the loading control. (**E**) RT-qPCR analysis of *CELSR2* mRNA expression using primers targeting the N-terminal region of *CELSR2.* Expression was normalized to *GAPDH* and is presented as relative quantification (RQ). (**F**) ChIP-seq tracks showing ZEB1 occupancy at the *CELSR2, VIM,* and *CDH1* loci in MDA-MB-231 cells (GSE89203). Signal tracks represent normalized read density, and ZEB1 binding peaks are highlighted in red. The position of the ATG start codon within exon 1 is indicated. (**G**) t-distributed stochastic neighbor embedding (t-SNE) visualization of single-cell RNA sequencing data from TNBC tumors (GSE161529), showing *CELSR2* (top) and *ZEB1* (bottom) expression projected onto the same embedding. (**H**) RT-qPCR analysis of *CELSR2, VCAM1, PDGFRA, EPCAM,* and *CDH1* expression in MDA-MB-468 cells transfected with control (siCtrl) or *CELSR2*-targeting siRNAs for 48 h. Gene expression was normalized to *GAPDH* and is presented as relative quantification (RQ). (**I**) RT-qPCR analysis of *CELSR2, CDH1, KRT18, KRT19, EPCAM, VIM, VCAM1,* and *PDGFRA* expression in MDA-MB-231 cells stably transduced with mCherry (mCh) or mCh+CELSR2 lentivirus. Gene expression was normalized to GAPDH and is presented as relative quantification (RQ). (**J**) *CELSR2* mRNA expression in circulating tumor cells (CTCs) (n=16) and primary tumors (n=12) from the BC cohort GSE111842. Expression values are shown as normalized log₂-transformed values. Box plots indicate the median and interquartile range. Statistical significance was assessed using a two-sided Wilcoxon rank-sum test. (**K**) *CELSR2* mRNA expression in primary (n = 44) and metastatic (n = 79) BC tumors from the GSE209998 cohort. Expression values are shown as normalized log₂-transformed values. Box plots indicate the median and interquartile range. Statistical significance was assessed using a two-sided Wilcoxon rank-sum test.

To further investigate the mechanisms underlying CELSR2 downregulation during EMT, we analyzed publicly available ChIP-seq datasets from the mesenchymal TNBC and CELSR2-negative cell line MDA-MB-231. ZEB1, a key EMT-associated transcription factor, was found to occupy regulatory regions within the *CELSR2* locus (**Fig. 2F**), supporting a potential role for ZEB1 in *CELSR2* regulation during EMT. Our analyses also identified ZEB1 binding at the *VIM* promoter whereas ZEB1 occupancy at both the *CDH1* (encoding E-Cadherin) and *CELSR2* loci was detected within the gene body. Given that *CDH1* expression is known to be repressed by ZEB1 during EMT progression^29^, these observations support a potential contribution of ZEB1 to *CELSR2* transcriptional repression. In agreement with these findings, single-cell RNA sequencing analyses revealed mutually exclusive expression patterns between *ZEB1* and *CELSR2* within TNBC tumor cells (**Fig. 2G**). Occupancy within the gene body of the *CELSR2* locus was also observed for another EMT-associated transcription factor, TWIST1, in the luminal breast cancer cell line MCF7 (**Fig. S3J**).

We next investigated whether CELSR2 loss could contribute to EMT. Silencing of CELSR2 in MDA-MB-468 cells using two independent siRNAs increased the expression of the mesenchymal markers *VCAM1* and *PDGFRA* (**Fig. 2H**), while epithelial markers remained largely preserved, with only a modest reduction in *CDH1* expression and no significant change in *EPCAM* expression (**Fig. 2H**), indicating that loss of CELSR2 promotes mesenchymal features and acquisition of a hybrid state. Conversely, ectopic expression of mCh-CELSR2 following lentiviral transduction in the mesenchymal MDA-MB-231 CELSR2-negative TNBC cells induced re-expression of the epithelial marker *KRT19* and downregulation of mesenchymal markers *VCAM1* and *PDGFRA* (**Fig. 2I**), supporting a role for CELSR2 in promoting epithelial features. CELSR2 overexpression in MDA-MB-231 cells was confirmed by fluorescence microscopy and western blot analysis (**Fig. S4A, B**). Consistently, ectopic expression of mCh-CELSR2 following lentiviral transduction of the mesenchymal MGT19 murine TNBC cells reduced the expression of several mesenchymal markers, including *Vimentin*, *Zeb1*, *Twist1*, and *Pdgfrb,* while Epcam and Cdh1 showed a non-significant trend toward increased expression (**Fig. S4C**). CELSR2 overexpression in MGT19 cells was confirmed by fluorescence microscopy and western blot analysis (**Fig. S4D, E**).

Consistent with the association between low CELSR2 expression and mesenchymal features, *in silico* analyses using bulk RNA-seq data from a publicly available breast cancer cohort (GSE111842) revealed reduced CELSR2 expression in circulating tumor cells (CTCs), which are frequently characterized by hybrid EMT states^9,30^ (**Fig. 2J**). Interestingly, analyses of metastatic lesions from another publicly available breast cancer cohort (GSE209998) showed re-expression of CELSR2 to levels comparable to those observed in primary tumors (**Fig. 2K**) consistent with a potential role for CELSR2 in epithelial reprogramming during MET. Altogether, these findings strongly support a role for CELSR2 in epithelial-mesenchymal plasticity in TNBC cells and suggest that variable CELSR2 expression levels may contribute to the phenotypic intratumoral heterogeneity observed in TNBC tumors.

### CELSR2 loss impairs cell-matrix adhesion while preserving epithelial cell-cell junctions

We next investigated whether CELSR2 loss induces changes associated with EMT such as loss of cell polarity and cell adhesion^6^. Despite their epithelial phenotype, MDA-MB-468 cells display poorly organized cell-cell junctions, making reliable assessment of junctional architecture by immunofluorescence difficult. We therefore used MCF7 cells, a luminal breast cancer cell line that forms well-organized epithelial cell-cell junctions, to evaluate the impact of CELSR2 depletion on junctional organization. We examined E-Cadherin and ZO-1 localization, as readouts of adherens and tight junction organization, respectively. Confocal Z-stack analyses showed that, despite efficient CELSR2 depletion, both E-Cadherin and ZO-1 retained a prominent localization at cell-cell contacts (**Fig. 3A**), with ZO-1 restricted to the apical junctional region, whereas E-cadherin and CELSR2 localized more basolaterally. These findings indicate that CELSR2 loss does not disrupt the overall organization of epithelial cell-cell junctions. Notably, some models of hybrid epithelial–mesenchymal states retain epithelial cell-cell junctions despite the acquisition of mesenchymal traits^31^.

**Fig. 3.**
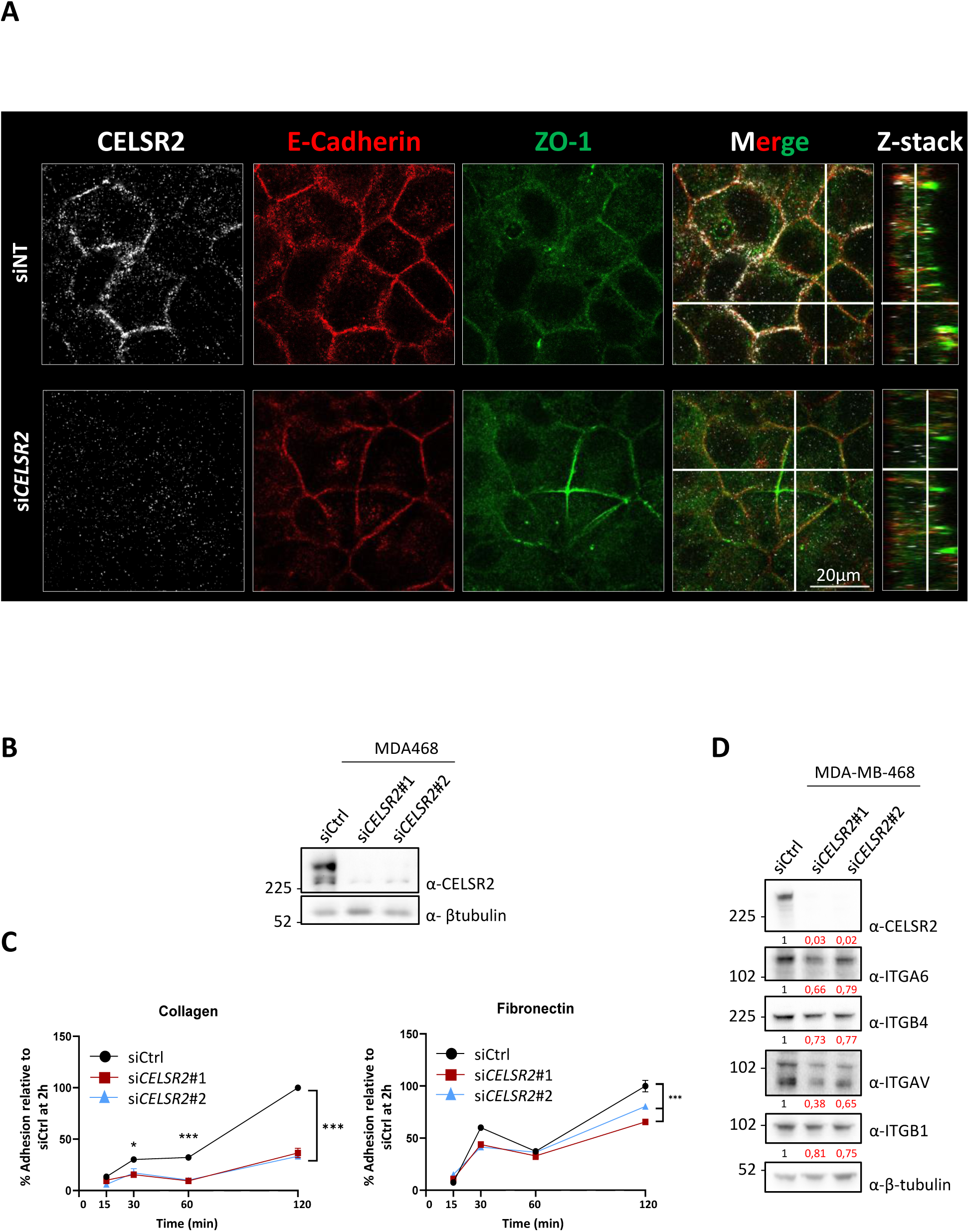
CELSR2 loss impairs cell–matrix adhesion while preserving epithelial cell–cell junctions. (**A**) Immunofluorescence analysis of E-cadherin, ZO-1, and CELSR2 expression in MCF-7 cells transfected with control (siCtrl) or CELSR2-targeting siRNA (si*CELSR2*#2). Representative XY images and orthogonal views from Z-stack acquisitions are shown. Scale bars, 20 μm. (**B**) Western blot analysis confirming *CELSR2* knockdown in MDA-MB-468 cells transfected with control (siCtrl) or *CELSR2*-targeting siRNAs. (**C**) Adhesion assay of MDA-MB-468 cells transfected with siCtrl or CELSR2-targeting siRNAs for 48 h and seeded onto collagen- (left) or fibronectin-coated (right) plates (10 μg/mL). Cell adhesion was quantified by crystal violet staining after 15 and 30 min, and 1 and 2 h of incubation. Results are expressed as the percentage of adhesion relative to the siCtrl condition at 2 h. Data are representative experiment from three independent experiments. (**D**) Western blot analysis of ITGA6, ITGB4, ITGAV, ITGB1, CELSR2, and β-tubulin expression in MDA-MB-468 cells transfected with siCtrl or CELSR2-targeting siRNAs. β-Tubulin served as the loading control.

Cell adhesion of silenced CELSR2 MDA-MB-468 cells was next assessed on collagen- and fibronectin-coated substrates. Efficient *CELSR2* silencing was confirmed by western blot analysis (**Fig. 3B**). *CELSR2* silencing significantly reduced cell adhesion to collagen at 30 min, 1 h, and 2 h after cell seeding. Adhesion to fibronectin was also decreased and reached statistical significance after 2 h (**Fig. 3C**). Representative images of adherent cells on collagen- and fibronectin-coated substrates at the indicated time points show reduced attachment of CELSR2-silenced cells compared with control cells (**Fig. S5**). To further investigate the effects of CELSR2 loss on cell-matrix adhesion, we analyzed the expression of the α6β4 integrin complex, a key mediator of epithelial cell-matrix adhesion and hemidesmosome organization^32^, and the αvβ1 integrin complex, a receptor for fibronectin and other RGD-containing extracellular matrix proteins^33^. *CELSR2* silencing was associated with reduced expression of ITGA6 and ITGB4, as well as ITGAV and ITGB1, corresponding to the α6β4 and αvβ1 integrin complexes, respectively (**Fig. 3D**). These observations further support a role for CELSR2 in regulating cell-matrix adhesion in TNBC cells. Taken together, these findings indicate that CELSR2 loss impairs cell-matrix adhesion while preserving epithelial cell-cell junctions.

### Spatial heterogeneity of *CELSR2* expression in TNBC tumors

Invasive tumor fronts are frequently enriched in EMT-associated programs^6^, and are increasingly recognized to correspond to hybrid EMT states^34,35^. As CELSR2 loss promotes the acquisition of a hybrid EMT phenotype, we sought to determine its expression at the invasive tumor front using publicly available spatial transcriptomic data from 94 TNBC patient samples^26^. Using the Hematoxylin and eosin (H&E) based tissue compartment annotations provided by Wang et al^26^., we identified a dominant class for each spot including tumor, stromal, immune, vascular, adipose, and ductal regions in a representative spatial transcriptomic tissue section (**Fig. 4A**). Projection of tumor marker expression intensity onto the same tissue section was used to estimate the tumor cell content of each spatial transcriptomic spot, with each spot representing approximately 15–20 cells (**Fig. 4B**). *CELSR2* expression was then mapped across the tissue section and was predominantly detected within tumor regions (**Fig. 4C**), consistent with findings reported in **Fig. 1F**. In agreement with its association with epithelial features, *CELSR2*-positive spots spatially co-localized with epithelial markers such as *CDH1*, whereas limited overlap was observed with mesenchymal markers such as *ZEB1* and *TWIST1* (**Fig. S6A, B**). To assess *CELSR2* spatial distribution within the tumor compartment, tumor core and invasive peripheral regions were defined as described in the Materials and Methods (**Fig. 4D**). To further validate the definition of tumor core and invasive peripheral regions, spatial mapping of epithelial and mesenchymal markers showed enrichment of *CDH1*-positive and ZEB1-negative spots within central tumor areas (**Fig. S6C, D**). Notably, *CELSR2*-positive spots were mainly detected in the tumor core, whereas invasive peripheral regions displayed fewer *CELSR2*-positive spots (**Fig. 4E**). Quantification across 94 TNBC patient samples confirmed a significantly higher proportion of *CELSR2*-positive spots in tumor core regions compared with invasive peripheral areas (**Fig. 4F**, **S6E**). Together, these findings indicate that CELSR2 expression is higher in the epithelial tumor core and reduced in invasive tumor regions, further supporting the association between low CELSR2 expression and a hybrid EMT phenotype, and suggesting that CELSR2 loss occurs preferentially at the invasive front.

**Fig. 4.**
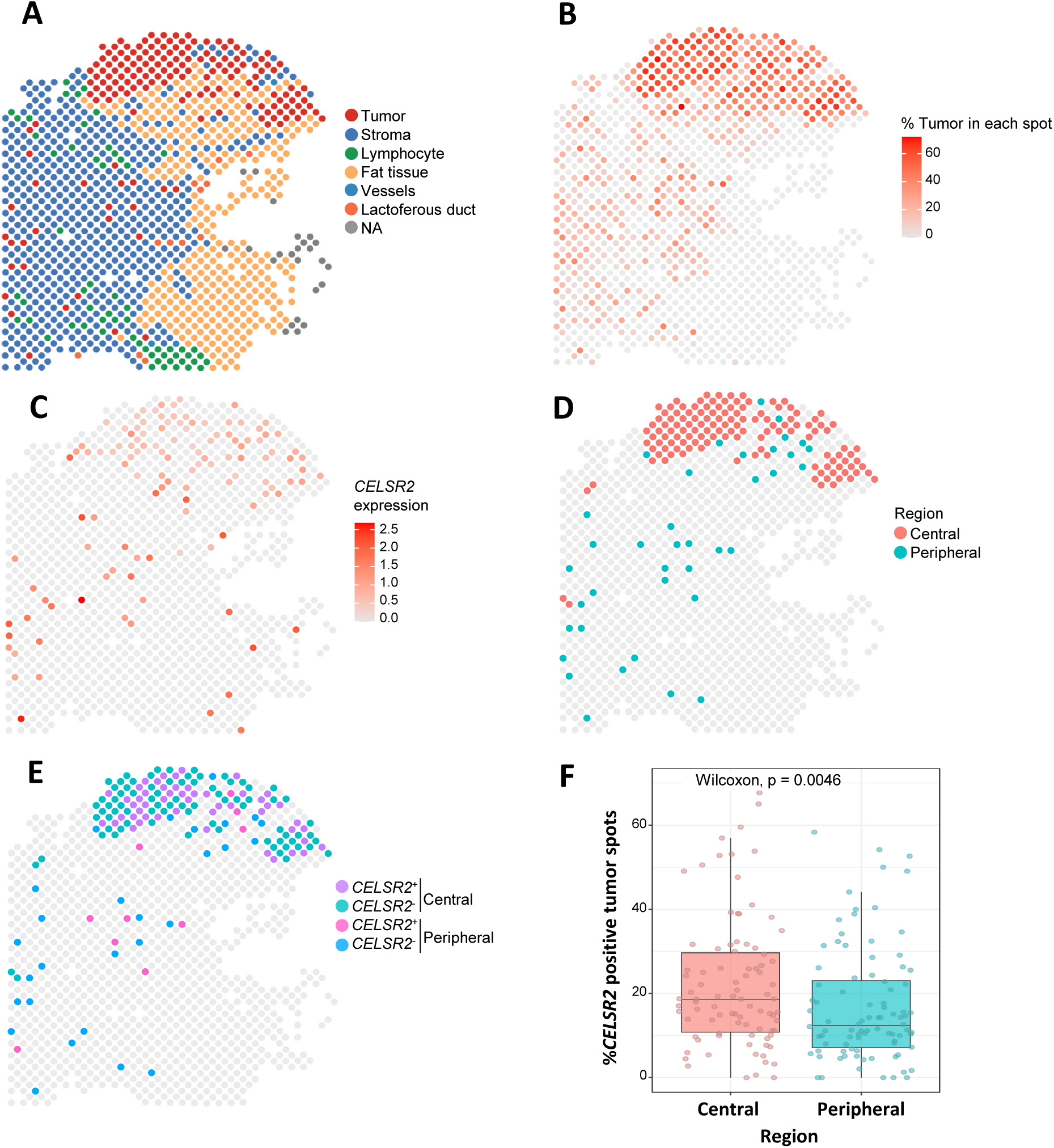
*CELSR2* expression is enriched in tumor core regions and reduced at the invasive front of TNBC tumors. (**A**) Representative H&E-based tissue compartment annotations showing tumor (red), stroma (blue), lymphocytes (green), blood vessels (blue), fat tissue (yellow), and lactiferous ducts (orange). (**B**) Spatial tumor purity map showing the percentage of tumor cells within each spot. Color intensity reflects the relative proportion of tumor cells within each spatial location. (**C**) Spatial distribution of *CELSR2* expression across the same tumor section. Expression values are shown as log-normalized transcript counts. (**D**) Spatial classification of tumor spots into central and peripheral regions using a *k*-nearest neighbor-based edge detection algorithm. Tumor spots were classified as peripheral when ≥25% of neighboring spots were non-tumoral and as central otherwise. (**E**) Distribution of *CELSR2*-positive and *CELSR2*-negative tumor spots in central and peripheral tumor regions. (**F**) Quantification of the proportion of *CELSR2*-positive tumor spots in central and peripheral regions across the TNBC cohort. Each point represents one patient. Statistical significance was assessed using a paired Wilcoxon signed-rank test (p = 0.0046).

### High CELSR2 expression promotes tumor growth and cell proliferation in TNBC, whereas low expression is associated with increased migratory properties

Given that CELSR2 expression is enriched in tumor core regions and reduced at the invasive front, we next investigated its role in TNBC progression using epithelial and mesenchymal TNBC cell lines with different CELSR2 expression levels. Consistent with their respective epithelial and mesenchymal states, MDA-MB-468 cells express higher levels of CELSR2 and E-Cadherin, whereas MDA-MB-231 cells have lower CELSR2 expression associated with high Vimentin expression (**Fig. 5A**). We first investigated the contribution of CELSR2 to tumor growth using stably silenced CELSR2 MDA-MB-468 cells generated by two independent lentiviral shRNAs (**Fig. 5B, C**). Orthotopic xenografts were then performed by injecting control or CELSR2-silenced MDA-MB-468 cells into the mammary fat pad of *NSG* mice, followed by assessment of primary tumor growth (**Fig. 5D**). Weekly monitoring revealed significantly reduced tumor volumes in mice injected with shCELSR2 cells compared with controls (**Fig. 5E**), whereas no significant difference in mouse body weight was observed during the experiment (**Fig. 5F**). Tumors were harvested and weighed at day 28. *CELSR2* silencing significantly reduced primary tumor weight compared with controls (**Fig. 5G, H**). Consistent with previous reports demonstrating that CELSR2 promotes tumor growth *in vivo* in glioma^16^, our results indicate that CELSR2 promotes primary tumor growth in TNBC.

**Fig. 5.**
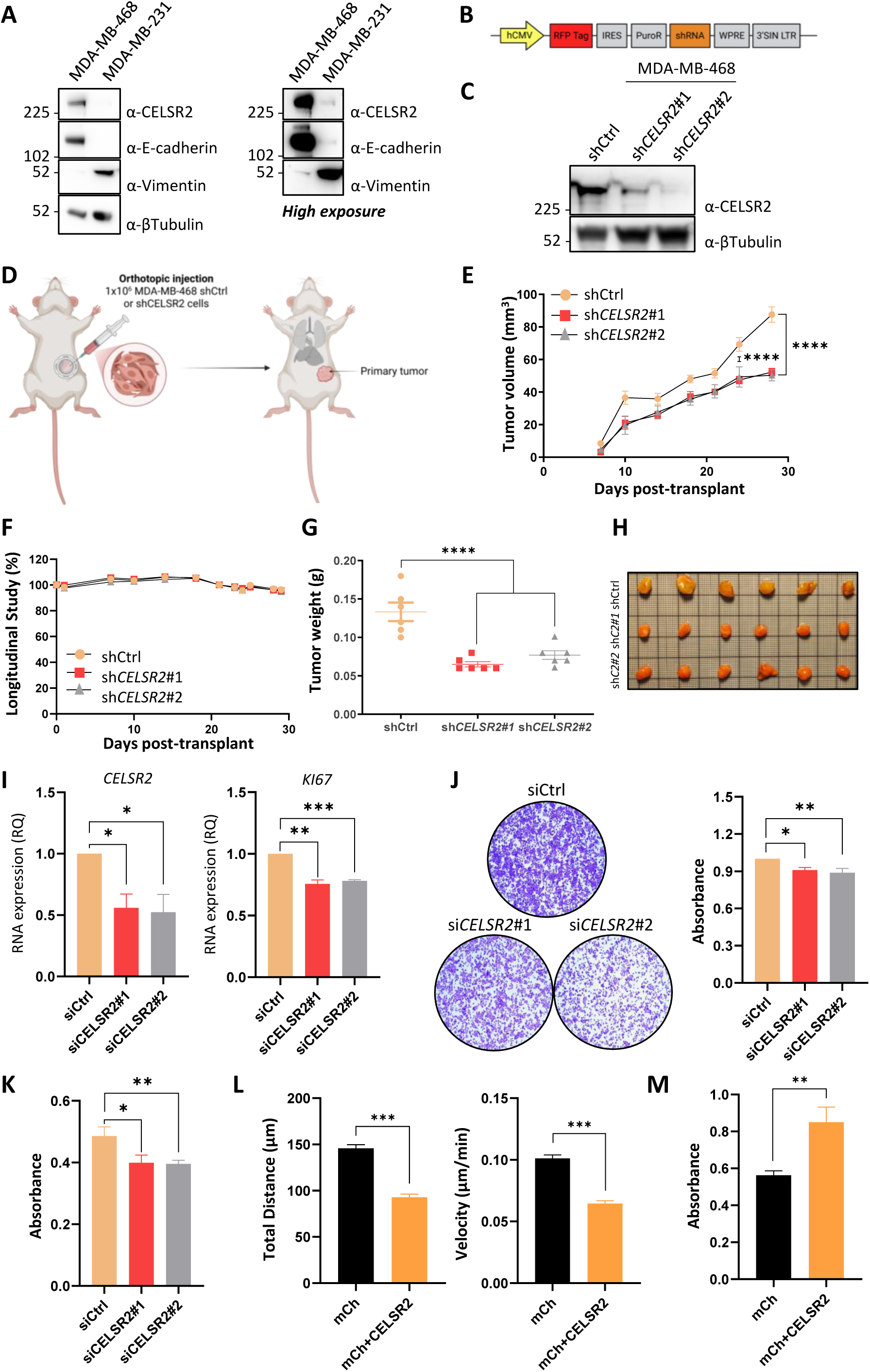
High CELSR2 expression promotes tumor growth and proliferation in TNBC cells whereas low expression is associated with increased cell migratory properties. (**A**) Western blot analysis of CELSR2, E-cadherin, Vimentin, and β-tubulin expression in MDA-MB-468 and MDA-MB-231 cells. β-Tubulin served as the loading control. (**B**) Schematic representation of the shRNA constructs used to knock down CELSR2. (**C**) Western blot analysis confirming *CELSR2* knockdown in MDA-MB-468 cells transduced with control (shCtrl) or *CELSR2*-targeting shRNAs. (**D**) Schematic representation of the orthotopic xenograft model. MDA-MB-468 cells (1 × 10⁶ cells/mouse; n = 6 mice per group) expressing control (shCtrl) or *CELSR2*-targeting shRNAs were orthotopically injected into mice. (**E**, **F**) Dynamical measurement of tumor volume (**E**) and body weight monitoring (**F**) of mice bearing shCtrl or shCELSR2 xenografts. (**G**, **H**) Quantification of primary tumor weight at the experimental endpoint (**G**) and representative excised tumors from each experimental group (**H**). (**I**) RT-qPCR analysis of *CELSR2* and *KI67* expression in MDA-MB-468 cells transfected with siCtrl or *CELSR2*-targeting siRNAs. Gene expression was normalized to *GAPDH* and is presented as relative quantification (RQ). (**J**) Crystal violet staining (left) and corresponding absorbance quantification (right) of MDA-MB-468 cells transfected with siCtrl or *CELSR2*-targeting siRNAs. (**K**) Cell proliferation measured by CCK-8 assay in MDA-MB-468 cells transfected with siCtrl or *CELSR2*-targeting siRNAs. (**L**) Quantification of total migration distance (μm) and mean migration velocity (μm/min) of MDA-MB-231 cells stably expressing mCherry (mCh) or mCh-CELSR2. Cell tracking was performed using ImageJ. (**M**) Cell proliferation measured by CCK-8 assay in MDA-MB-231 cells stably expressing mCherry (mCh) or mCh-CELSR2. Data represent the mean ± SEM of 3 independent experiments. The in vivo experiment shown is representative of two independent experiments. Statistical significance was determined using two-sided Student’s *t*-tests for in vitro experiments and two-way ANOVA followed by Tukey’s multiple-comparisons test for the in vivo experiment (\**P* < 0.05, ** *P* < 0.01, \*\*\**P* < 0.001, **** *P* < 0.0001).

To determine whether reduced tumor growth following *CELSR2* silencing could be associated with altered cell proliferation, we next assessed expression of the proliferation marker Ki-67 in MDA-MB-468 cells. Ki-67 expression was significantly reduced following *CELSR2* silencing (**Fig. 5I**). Consistent with these findings, *CELSR2* silencing reduced cell number, as assessed by crystal violet staining, and decreased the CCK-8 signal (**Fig. 5J, K**). Together with reduced Ki-67 expression, these findings support impaired proliferative capacity following *CELSR2* silencing. Similarly, *CELSR2* silencing in the epithelial-like MGT14 cell line reduced absorbance following crystal violet staining (**Fig. S7A**). These findings are consistent with previous studies reporting a pro-proliferative role for CELSR2 in breast cancer^18^, hepatocellular carcinoma^17^, and glioma^16^. Altogether, these results indicate that CELSR2 promotes TNBC tumor growth and supports cell proliferation of TNBC cells.

To gain insight into the contribution of CELSR2 to cellular traits associated with metastatic dissemination, we examined the effects of stable CELSR2 overexpression in mesenchymal MDA-MB-231 TNBC cells with low CELSR2 levels. Single-cell tracking analyses revealed that CELSR2 overexpression significantly reduced both total migration distance and cell velocity compared with controls (**Fig. 5L**). These results suggest that reduced CELSR2 expression leads to the acquisition of migratory properties, consistent with the acquisition of hybrid epithelial–mesenchymal features described above.

Because metastatic colonization and outgrowth require reacquisition of proliferative capacities following dissemination, we next investigated whether CELSR2 expression could promote growth-related properties in mesenchymal TNBC cells. CELSR2 overexpression significantly increased metabolic activity in MDA-MB-231 cells, as assessed by CCK-8 assays (**Fig. 5M**). Similar results were obtained in mesenchymal MGT19 cells stably expressing CELSR2 (**Fig. S7B**). These findings indicate that CELSR2 overexpression promotes proliferative properties in mesenchymal TNBC cells.

Taken together, these data indicate that reduced CELSR2 expression favors a migratory hybrid EMT state, whereas high CELSR2 expression is associated with proliferative properties and epithelial features. These observations are consistent with the spatial distribution of CELSR2 within TNBC tumors, where CELSR2^high^ tumor cells are enriched in the proliferative epithelial tumor core and CELSR2^low^ tumor cells preferentially detected at the migratory invasive front. Collectively, these findings support the notion that heterogeneous CELSR2 expression contributes to distinct cell states associated with different tumoral properties during TNBC progression.

### CELSR2 promotes a CREB-associated OXPHOS program in TNBC cells

We next performed proteomic analyses of MDA-MB-468 cells following *CELSR2* silencing. Mass spectrometry revealed a reduced abundance of several proteins encoded by CREB target genes upon *CELSR2* silencing, as shown in the heatmap in **Fig. S8A**. Consistently, similar results were obtained in MCF7 cells following *CELSR2* silencing (**Fig. S8B**). In line with these observations, recent work identified CELSR2 as an upstream activator of Gαs signaling, promoting cAMP production and suggesting that CELSR2 may regulate CREB activity^27^. To determine whether CELSR2 activates CREB, HEK293T cells were transiently co-transfected with a CRE-responsive Firefly luciferase reporter, a Renilla luciferase control plasmid, and either GFP or CELSR2-GFP expression vectors (**Fig. 6A**) whose expression was confirmed by fluorescence microscopy (**Fig. S9A**) and western blot (**Fig. S9B**). CELSR2 expression significantly increased CREB-dependent luciferase activity (**Fig. 6A**). Western blot analyses showed increased CREB phosphorylation at Ser_133_ in CELSR2-GFP-transfected cells compared with GFP and CELSR1-GFP controls (**Fig. 6B**, **Fig. S9B**), further supporting activation of CREB signaling downstream of CELSR2.

**Fig. 6.**
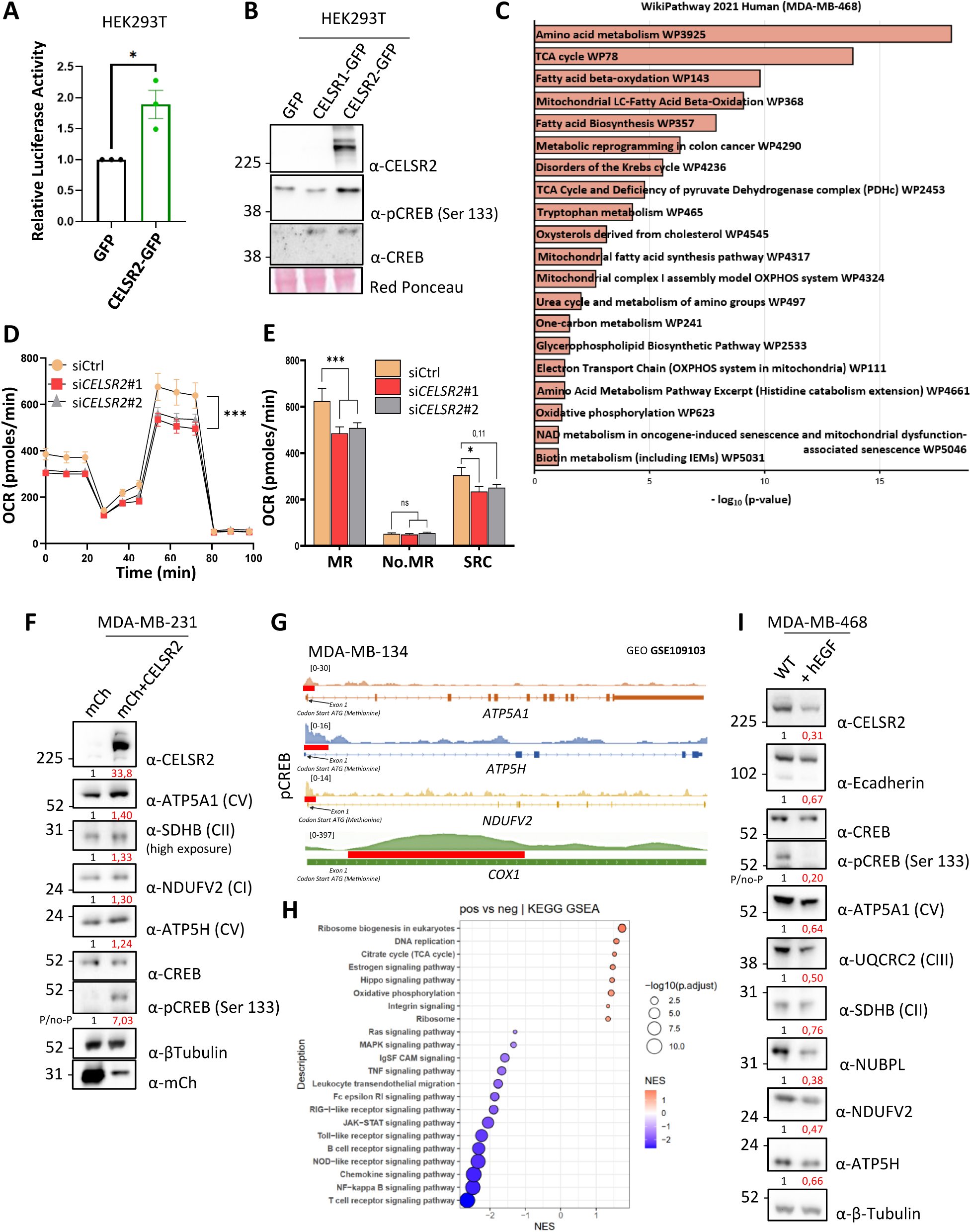
CELSR2 promotes CREB activation and OXPHOS in TNBC cells. (**A**) CRE luciferase reporter activity in HEK293T cells transfected with GFP or GFP-CELSR2 expression plasmids. Luciferase activity was normalized to the mean value of GFP-transfected cells. (**B**) Western blot of GFP, CELSR1-GFP and CELSR2-GFP HEK293T cells showing the expression of CELSR2, pCREB (Ser133) and CREB. Ponceau S staining served as the loading control. (**C**) WikiPathways 2021 Human enrichment analysis of proteins downregulated following CELSR2 silencing in MDA-MB-468 cells. The 20 highest-ranking pathways are displayed, with bar length representing −log10(P value). (**D**) Oxygen consumption rate (OCR) was measured in MDA-MB-468 cells transfected with siCtrl or *CELSR2*-targeting siRNAs using the Seahorse Mito Stress Test. OCR was recorded under basal conditions and following sequential injections of oligomycin, carbonyl cyanide-*p*-trifluoromethoxyphenylhydrazone (FCCP), and rotenone plus antimycin A. (**E**) Maximal respiration (MR), non-mitochondrial respiration (No.MR), and spare respiratory capacity (SRC) derived from Seahorse Mito Stress Test analysis of MDA-MB-468 cells transfected with siCtrl or *CELSR2*-targeting siRNAs. (**F**) Western blot analysis of CELSR2, ATP5A1, SDHB, NDUFV2, ATP5H, mCherry, and β-tubulin expression in MDA-MB-231 cells expressing mCherry (mCh) or mCh+CELSR2. β-Tubulin served as the loading control. The mCh-CELSR2 plasmid contains a T2A self-cleaving peptide, resulting in the independent expression of mCh and CELSR2. (**G**) ChIP-seq tracks showing pCREB occupancy at the *ATP5A1*, *ATP5H*, *NDUFV2*, and *COX1* loci in MDA-MB-134 cells (GSE109103). Signal tracks represent normalized read density, and pCREB binding peaks are highlighted in red. The position of the ATG start codon within exon 1 is indicated. (**H**) KEGG pathway enrichment analysis comparing *CELSR2*-positive and *CELSR2*-negative tumor cells from TNBC single-cell RNA sequencing data ^62^(GSE176078). Positive normalized enrichment scores (NES) indicate pathways enriched in *CELSR2*-positive cells, whereas negative NES values indicate pathways enriched in *CELSR2*-negative cells. (**I**) Western blot analysis of CELSR2, E-cadherin, CREB, phospho-CREB (Ser133), ATP5A1, UQCRC2, SDHB, NUBPL, NDUFV2, ATP5H, and β-tubulin expression in untreated (WT) and hEGF-treated MDA-MB-468 cells. β-Tubulin served as the loading control. The P/no-P ratio corresponds to the abundance of phosphorylated CREB relative to total CREB.

To gain insight into the biological processes regulated by CELSR2 in TNBC cells, Gene Ontology enrichment analyses were performed using the proteomic dataset from CELSR2-silenced MDA-MB-468 cells. We identified oxidative phosphorylation (OXPHOS) along with several metabolic pathways fueling mitochondrial respiration, including amino acid metabolism, fatty acid β-oxidation, and the TCA cycle (**Fig. 6C**). We next assessed OXPHOS activity in CELSR2-silenced cells using Seahorse analyses. Efficient CELSR2 downregulation was confirmed by western blot (**Fig. S9C**). Oxygen consumption rate (OCR), a functional readout of OXPHOS, was measured in control and CELSR2-silenced MDA-MB-468 cells (**Fig. 6D**). *CELSR2* silencing reduced basal respiration, maximal respiration (MR), and spare respiratory capacity (SRC), indicating impaired mitochondrial oxidative metabolism upon *CELSR2* silencing (**Fig. 6E**).

We also investigated whether CELSR2 expression was sufficient to promote CREB signaling and oxidative phosphorylation in TNBC cells. Stable CELSR2 overexpression in MDA-MB-231 cells increased CREB phosphorylation at Ser_133_ compared with control cells, indicating activation of CREB signaling downstream of CELSR2 (**Fig. 6F**). In parallel, CELSR2 overexpression increased the levels of multiple OXPHOS proteins in MDA-MB-231 cells (**Fig. 6F**). Similar increases in CREB phosphorylation and OXPHOS protein expression were observed in mesenchymal MGT19 cells following CELSR2 expression (**Fig. S9D, E**). Together, these findings support a role for CELSR2 in maintaining OXPHOS in epithelial TNBC cells, in addition to its ability to activate CREB signaling, two pathways known to support tumor cell proliferation and tumor growth^36–39^. To determine whether CREB could directly regulate oxidative phosphorylation-related genes downstream of CELSR2, publicly available CREB ChIP-seq datasets were analyzed in MDA-MB-134 cells (GSE109103). Promoter-proximal pCREB1 peaks were observed at the OXPHOS-related genes *ATP5A1, ATP5H, NDUFV2,* and *COX1* (**Fig. 6G**), supporting their identification as candidate pCREB1-bound genes. These observations, together with previous reports identifying direct regulation of mitochondrial and OXPHOS genes by CREB signaling ^40–43^, support a role for CREB in regulating the OXPHOS program downstream of CELSR2. Finally, KEGG gene set enrichment analyses using an independent TNBC single-cell RNA-seq cohort revealed that CELSR2-positive tumor cells were enriched in metabolic and proliferative pathways, including oxidative phosphorylation, ribosome biogenesis, and DNA replication (**Fig. 6H**). Together, these findings identify a CREB-associated OXPHOS program downstream of CELSR2 that provides a potential mechanistic basis for the increased cell proliferation and primary tumor growth promoted by CELSR2 (**Fig. 5**).

Given that CELSR2 regulates epithelial-mesenchymal plasticity and promotes OXPHOS, we next explored a putative link between CELSR2-associated metabolic changes and epithelial-mesenchymal features. EMT was induced in MDA-MB-468 cells using hEGF treatment, as previously described (**Fig. 2**), leading to reduced CELSR2 expression (**Fig. 6I**). This reduction was accompanied by decreased CREB phosphorylation and lower expression of OXPHOS proteins (**Fig. 6I**), indicating that the transition toward a mesenchymal state is associated with attenuation of the CELSR2-CREB-OXPHOS axis. These observations suggest that the effect of CELSR2 on cellular metabolism may be linked, at least in part, to its regulation of epithelial–mesenchymal plasticity. These findings are consistent with previous reports describing an inverse relationship between OXPHOS activity and EMT programs^44^.

These results identify CELSR2 as a regulator of a CREB-associated oxidative phosphorylation program linked to epithelial tumor cell states. This program is maintained in CELSR2^high^ epithelial TNBC cells, diminished following EMT induction, and enhanced upon CELSR2 expression in mesenchymal TNBC cells. These results provide a potential mechanistic link between CELSR2 expression, oxidative metabolism, and the proliferative properties that support tumor growth. Collectively, our data support a model in which heterogeneous CELSR2 expression contributes to the emergence of distinct epithelial-mesenchymal states with different metabolic and proliferative properties, thereby contributing to intratumoral heterogeneity in TNBC.

## Discussion

TNBC is characterized by marked intratumoral heterogeneity, which contributes to tumor progression, therapeutic resistance, and metastatic dissemination. This heterogeneity is driven by multiple factors, including genetic, immune, microenvironmental, metabolic, and cell-state differences. Epithelial-mesenchymal plasticity represents one component of this heterogeneity and contributes to the emergence of tumor cell populations with distinct phenotypic and functional properties. Here, we identify CELSR2 as a regulator of epithelial–mesenchymal plasticity in TNBC. We show that CELSR2 expression promotes epithelial traits together with a CREB-associated oxidative phosphorylation program, proliferative capacity and primary tumor growth *in vivo*. Conversely, CELSR2 loss leads to the acquisition of a hybrid EMT state. Consistent with these observations, spatial transcriptomic analyses revealed a preferential localization of CELSR2^high^ tumor cells in the tumor core and of CELSR2^low^ tumor cells at the invasive front.

Interestingly, although high *CELSR2* expression is associated with shorter MFS in TNBC (**Fig. 1**) and contributes to TNBC progression *in vivo* (**Fig. 5**), CELSR2 was not overexpressed in TNBC compared with tumor-adjacent breast tissues (AT) or normal breast tissues from healthy people (**Fig. S10**). This apparent paradox is likely explained by the pronounced epithelial-mesenchymal heterogeneity that characterizes TNBC, both within individual tumors^5^ and across patients^45^. Because CELSR2 expression is largely restricted to epithelial tumor cells (**Fig. 1**, **Fig. 2**), its overall expression in bulk TNBC samples is diluted by the presence of CELSR2^low^ mesenchymal tumor cells.

A second apparent paradox is the association between high CELSR2 expression and poor MFS despite its association with epithelial cell states. One explanation comes from our *in vivo* data which demonstrate that CELSR2 promotes primary tumor growth. Since primary tumor growth is a major determinant of metastatic dissemination^46^, CELSR2^high^ proliferative tumor cells may therefore increase the risk of metastatic progression. A second possibility is that high CELSR2 levels in primary tumors are subsequently found in metastatic lesions, where CELSR2 could contribute to MET-dependent processes of metastatic colonization and outgrowth^8,9,47^. This hypothesis is supported by the re-expression of CELSR2 in metastatic lesions (**Fig. 2**), together with its ability to increase the proliferative capacity of mesenchymal TNBC cells (**Fig. 5**, **Fig. S7**), suggesting that its re-expression may promote metastatic outgrowth.

Whereas CELSR2^high^ tumor cells were enriched in the tumor core, consistent with the growth-promoting effects^48^ of CELSR2, CELSR2^low^ tumor cells preferentially localized to the invasive front (**Fig. 4**), a region associated with tumor cell invasion and dissemination^9^. Consistently, reduced CELSR2 expression promoted the acquisition of hybrid epithelial-mesenchymal characteristics (**Fig. 2**), a cellular state known to promote metastatic dissemination^31^. Further supporting a link between reduced CELSR2 expression and tumor cell dissemination, we found decreased CELSR2 expression in circulating tumor cells (**Fig. 2**). Together, these observations suggest that TNBC progression relies on the complementary functions of both CELSR2^high^ and CELSR2^low^ tumor cell populations.

The acquisition of migratory and invasive properties during the establishment of hybrid EMT states is accompanied by extensive remodeling of cell-matrix interactions, including reduced expression of α6β4 and increased expression of integrins that mediate adhesion to collagen I and fibronectin^49,50^. Consistently, *CELSR2* silencing reduced ITGA6/ITGB4 expression (**Fig. 3**). However, CELSR2 depletion also impaired adhesion to collagen I and fibronectin. These findings suggest that the cell-matrix adhesion program associated with CELSR2 loss does not fully recapitulate the canonical integrin remodeling described during EMT. This may reflect the context-dependent nature of EMT^51^, with distinct transcriptional programs generating phenotypically diverse hybrid EMT states. Alternatively, given the established role of adhesion GPCRs in regulating cell-matrix interactions^52^, impaired extracellular matrix adhesion could also represent a direct consequence of CELSR2 loss, independent of its effects on epithelial–mesenchymal plasticity.

Epithelial-mesenchymal plasticity has been increasingly recognized as a regulator of metabolic reprogramming in cancer^53^. Across several cancer types, EMT has been associated with reduced OXPHOS and increased glycolytic metabolism^44^. Consistently, recent proteomic profiling identified four molecular clusters in TNBC, including one enriched in OXPHOS and fatty acid metabolism, and a distinct cluster characterized by a strong EMT signature^45^. Together with our observation that EMT is accompanied by a concomitant decrease in *CELSR2* expression and OXPHOS (**Fig. 6**), these findings suggest that intratumoral heterogeneous *CELSR2* expression may contribute to the emergence of distinct metabolic states through its regulation of epithelial-mesenchymal plasticity. This model predicts that CELSR2^low^ hybrid epithelial-mesenchymal tumor cells would display increased glycolytic activity compared with their CELSR2^high^ epithelial counterparts. It also raises the possibility that the increased OXPHOS observed following CELSR2 overexpression in mesenchymal TNBC cells (**Fig. 6**) reflects the reacquisition of epithelial features. Both possibilities warrant further investigation. Nevertheless, such a model is likely an oversimplification, as metabolic programs associated with EMT are highly context-dependent and OXPHOS has also been implicated in metastatic dissemination in TNBC^54,55^.

More broadly, CELSR2 belongs to the adhesion GPCR family and is a component of the WNT/planar cell polarity (PCP) pathway, both of which have emerged as important regulators of cancer progression^11,52^. In breast cancer, several adhesion GPCRs, including ADGRF5/GPR116 and ADGRG1/GPR56, together with WNT/PCP components such as VANGL1, VANGL2, and PRICKLE1, have been implicated in tumor aggressiveness and metastatic dissemination through overexpression^10,12,56–58^. Rather than uniformly high CELSR2 expression, our findings support a model in which intratumoral heterogeneity of CELSR2 expression contributes to the aggressive behavior of TNBC by generating spatially (**Fig. 4**) and functionally (**Fig. 5**) distinct tumor cell populations that could collectively enable primary tumor growth and metastatic dissemination to occur simultaneously.

This spatial organization suggests the potential of CELSR2 as a spatial biomarker in TNBC. Compared with other breast cancer subtypes, TNBC still lacks validated prognostic and predictive biomarkers to guide therapeutic decision-making^59^. Our findings raise the possibility that assessment of the spatial distribution of CELSR2, rather than assessment of its overall expression level, could provide additional information on tumor behavior. High CELSR2 expression within the tumor core may identify highly proliferative epithelial tumor cell populations, whereas reduced CELSR2 expression at the invasive front may delineate regions enriched in tumor cells with dissemination-associated features. Validation of these spatial patterns by immunohistochemistry in an independent TNBC cohort will be required to determine whether CELSR2 could ultimately have value as a spatial biomarker associated with metastatic risk or clinical outcome.

Beyond the spatial organization of tumor cell states, their dynamic interconversion further contributes to tumor progression. Epithelial-mesenchymal plasticity is increasingly recognized as a therapeutic vulnerability in TNBC^5^, as reversible transitions between epithelial and mesenchymal states allow tumor cells to adopt phenotypes that support distinct stages of disease progression. Our study suggests that CELSR2 may participate in these phenotypic transitions. Notably, CELSR2 overexpression in mesenchymal TNBC cells did not elicit a complete reversion to an epithelial state (**Fig. 2**, **Fig. S4**), consistent with accumulating evidence that metastatic cells often undergo only partial MET^60^. By retaining mesenchymal traits, these incompletely reverted cells are likely highly aggressive and may preserve the phenotypic plasticity required to re-engage mesenchymal programs, potentially facilitating subsequent rounds of metastatic dissemination^31,61^. Whether selective modulation of CELSR2 can be exploited to limit epithelial-mesenchymal plasticity remains to be determined. As with other epithelial regulators of epithelial-mesenchymal plasticity, therapeutic modulation of CELSR2 is likely to present important challenges. Therapeutic inhibition of CELSR2 could suppress primary tumor growth while simultaneously promoting tumor cell states associated with dissemination. In this context, selective elimination of CELSR2-expressing tumor cells using antibody-drug conjugates, rather than inhibition of *CELSR2* signaling may represent a more effective therapeutic strategy.

Some limitations of this study should be acknowledged. First, whether activation of OXPHOS is directly responsible for the growth-promoting effects of CELSR2 remains to be formally established. Second, while our findings suggest that CELSR2 may influence metastatic progression through its regulation of epithelial-mesenchymal plasticity, this possibility was not directly addressed *in vivo*. Because CELSR2 may contribute to epithelial reprogramming during metastatic progression, stable CELSR2 knockdown cells were only used to investigate primary tumor growth *in vivo* rather than metastatic dissemination, as persistent *CELSR2* silencing would prevent its re-expression that may occur during later stages of the metastatic cascade. An inducible CELSR2 knockdown model would help overcome this limitation by enabling temporal control of CELSR2 expression and allowing its role to be dissected at distinct stages of metastatic progression. Finally, although spatial transcriptomic analyses revealed distinct spatial distributions of CELSR2^high^ and CELSR2^low^ tumor cell populations within TNBC lesions, these observations will require validation at the protein level.

Together, these findings support a model in which heterogeneous CELSR2 expression contributes to the coexistence of spatially and functionally distinct tumor cell populations characterized by different epithelial-mesenchymal states and oxidative metabolic properties, thereby promoting intratumoral heterogeneity in TNBC. Future studies should further explore whether CELSR2 acts as a regulatory node that coordinates epithelial-mesenchymal plasticity and metabolic reprogramming during tumor progression. Determining whether the spatial organization of CELSR2 expression is associated with clinical outcome will be an important step toward translating these findings into clinical practice.

## Acknowledgements

We would like to thank people of CRCM platforms: Manon Richaud of the flow cytometry platform and M. Rodrigues of the microscopy and scientific imaging platform. We are grateful to the staff of the CRCM animal facility for taking care of the mouse colonies. We thank Evelyne Thi Thien Nguyen for her assistance in the sea horse experiments setup. *We thank the Canceropôle PACA*, IBISA (Infrastructures en Biologie Santé et Agronomie), and the Plan Cancer Equipement (#17CQ047-00) for continued support in the development of the TrGET preclinical assay platform. Proteomics analyses were done at the mass spectrometry facility of Marseille Proteomics supported by Institut Paoli-Calmettes, CRCM, IBISA, ITMO Cancer, Aix-Marseille University, Canceropôle PACA, the Provence-Alpes-Côte d’Azur Région and the Fonds Européen de Développement Régional (FEDER).

We would like to thank Ines Liebscher, Luca Lignitto, Alice Carrier, Franck Vandermoere, Rania Ghossoub and Charlotte Dessaux for their help and fruitful discussion. We thank Danelle Devenport for the Celsr1-GFP construct and Tadashi Uemura for the Celsr2-GFP construct.

## Funding sources

This work has been supported by Inserm, Institut Paoli-Calmettes, the Ligue Nationale Contre le Cancer (Label Ligue JP Borg 2022), the Institut Universitaire de France, and Canceropole PACA.

JPB is a scholar of the Institut Universitaire de France; JK received a fellowship from the Ligue Nationale Contre le Cancer; NIYS received a fellowship from Cancer and Immunlogy Institute of Aix Marseille University.

## Authors contribution

Conceptualization: AW, JPB. Methodology: JPB, AW, FM, JK, NIYS. Collection, analyses and/or interpretation of the data: JK, NIYS, AE, SA, LC, AD, RC, FB, FL, PF, EJ, RG.

## Funding

acquisition & administration: JPB, AW, AGC. Supervision: JPB, FM, AW. Writing - original draft: AW, JPB. Writing - finalize, review & editing: FM, AW, JPB, JK, AE. All authors reviewed the results, proofread the manuscript and approved the final version of the manuscript.

## Declaration of interests

The authors declare no conflicts of interest.

## Figure Legends

**Supplementary Figure S1. CELSR2 expression correlates with epithelial features and is reduced in mesenchymal TNBC mouse cell lines.** (**A**) Western blot analysis of CELSR2 (normal exposure, left ; high exposure, right), E-cadherin, N-cadherin, and Vimentin expression in MGT cell lines. MGT cell lines displaying the most epithelial-like and the most mesenchymal-like phenotypes are indicated in blue and purple, respectively. Ponceau S staining served as the loading control. (**B**) RT-qPCR analysis of *Celsr2* expression in MGT cell lines using primer pairs targeting the N-terminal (NTF, left) or C-terminal (CTF, right) region of *Celsr2*. Gene expression was normalized to *Gapdh* and is presented as relative expression. Data represent the mean ± SEM of 3 independent experiments. Statistical significance was determined using two-sided Student’s *t*-tests. *P* < 0.05.

**Supplementary Figure S2. Association between the EMT score and CELSR2 expression in TNBC tumors.** Association between the Taube et al’s EMT metagene score and *CELSR2* mRNA expression in TNBC tumors of the whole series (n = 747; top) and in each of the 14 data sets separately. The forest plots display the Geometric Mean Ratio (GMR) in the whole series (top; n = 747) and in each data set separately (middle), and the pooled GMR (diamond) at the bottom. Heterogeneity among datasets was quantified using Cochran’s Q test.

**Supplementary Figure S3. Regulation of *Celsr2* expression during EMT.** (**A**) Top: Experimental design for EMT induction in MGT20 and MGT14 epithelial TNBC cells. Cells were left untreated (WT) or stimulated with transforming growth factor-β (TGF-β; 10 ng/mL) on day 1, followed by a second treatment with TGF-β (5 ng/mL) 72 h later. Cells were harvested on day 5. (**B**) Representative images of untreated (WT) and TGF-β-treated MGT20 cells. (**C**) RT-qPCR analysis of epithelial (top) and mesenchymal (bottom) marker expression in untreated (WT) and TGF-β-treated MGT20 cells. Gene expression was normalized to *Gapdh* and is presented as relative quantification (RQ). (**D**) Western blot analysis of CELSR2, E-cadherin, N-cadherin, EPCAM, Vimentin, and β-tubulin expression in untreated (WT) and TGF-β-treated MGT20 cells. β-Tubulin served as the loading control. (**E**) RT-qPCR analysis of *Celsr2* expression using primer pairs targeting the N-terminal region of *Celsr2*. Gene expression was normalized to *Gapdh* and is presented as relative quantification (RQ). (**F**) Representative images of untreated (WT) and TGF-β-treated MGT14 cells. (**G**) RT-qPCR analysis of epithelial (top) and mesenchymal (bottom) marker expression in untreated (WT) and TGF-β-treated MGT14 cells. Gene expression was normalized to *Gapdh* and is presented as relative quantification (RQ). (**H**) Western blot analysis of CELSR2, E-cadherin, N-cadherin, EPCAM, Vimentin, and β-tubulin expression in untreated (WT) and TGF-β-treated MGT14 cells. β-Tubulin served as the loading control. (**I**) RT-qPCR analysis of *Celsr2* expression using primer pairs targeting the N-terminal (left) or C-terminal (right) region of *Celsr2*. Gene expression was normalized to *Gapdh* and is presented as relative quantification (RQ). (**J**) ChIP-seq tracks showing TWIST1 occupancy at the *CELSR2* locus in MCF-7 cells (GSE189826). Signal tracks represent normalized read density, and TWIST1 binding peak is highlighted in red. The position of the ATG start codon within exon 1 is indicated. Data represent the mean ± SEM of 3 independent experiments. Statistical significance was determined using two-sided Student’s *t*-tests (\**P* < 0.05, \*\**P* < 0.01, \*\*\**P* < 0.001).

**Supplementary Figure S4. Ectopic expression of CELSR2 in mesenchymal TNBC cell lines induces re-expression of epithelial markers.** (**A**, **B**) Representative images of MDA-MB-231 cells stably expressing mCherry (mCh) or mCh+CELSR2 (**A**) and corresponding western blot analysis (**B**). The mCh-CELSR2 construct contains a T2A self-cleaving peptide, resulting in the independent expression of mCherry and CELSR2. (**C**) RT-qPCR analysis of *Celsr2*, *Cdh1*, *Epcam*, *Vim*, *Zeb1*, *Zeb2*, *Twist1*, and *Pdgfrb* expression in MGT19 cells stably expressing mCherry (mCh) or mCh+CELSR2. Gene expression was normalized to *Gapdh.* Data are presented as 1 + log2FC, with the control value set to 1. (**D**, **E**) Representative images of MGT19 cells stably expressing mCherry (mCh) or mCh+CELSR2 (**D**) and corresponding western blot analysis (**E**). The mCh-CELSR2 construct contains a T2A self-cleaving peptide, resulting in the independent expression of mCherry and CELSR2.

**Supplementary Figure S5. CELSR2 contributes to cell–matrix adhesion.** Representative images of MDA-MB-468 cells transfected with siCtrl or *CELSR2-*targeting siRNAs, showing adherent cells after 1 or 2 h on collagen- (top) or fibronectin-coated (bottom) plates (10 μg/mL).

**Supplementary Figure S6. Spatial association of CELSR2 with epithelial and mesenchymal markers in TNBC tumors.** (**A**) Spatial co-expression of *CELSR2* with epithelial and mesenchymal markers. (**B**) Spatial co-expression of *CELSR2* and *ZEB1*. (**C**) Distribution of *ZEB1-*positive tumor spots in central and peripheral tumor regions. Each dot represents a tumor spot classified as positive or negative for *ZEB1* detection. (**D**) Distribution of *CDH1*-positive tumor spots in central and peripheral tumor regions. Each dot represents a tumor spot classified as positive or negative for *CDH1* detection. (**E**) Paired comparison of *CELSR2*-positive tumor spots between central and peripheral tumor regions. Line colors indicate the outcome of Fisher’s exact test comparing the proportion of positive and negative tumor spots between regions for each sample (red, higher in the central region; green, lower in the central region; gray, not significant). Box plots show the distribution of *CELSR2*-positive tumor spots across samples. Statistical significance was assessed using the Wilcoxon signed-rank test (*P* = 0.0092).

**Supplementary Figure S7. CELSR2 is involved in the proliferation of mouse TNBC cells.** (**A**) Crystal violet staining (right) and corresponding absorbance quantification (left) of MGT14 cells transfected with siCtrl or *Celsr2*-targeting siRNAs. (**B**) Cell proliferation assessed by CCK-8 assay in MGT19 cells stably expressing mCherry (mCh) or mCh+CELSR2. Data represent the mean ± SEM of 3 independent experiments. Statistical significance was determined using two-sided Student’s *t*-tests (\**P* < 0.05, \*\**P* < 0.01, \*\*\**P* < 0.001).

**Supplementary Figure S8. CELSR2 silencing alters CREB-associated proteins and mitochondrial metabolism in breast cancer cells.** (**A–B**) Heatmaps showing the relative abundance of CREB motif-associated targets and highly differential proteins in MDA-MB-468 (**A**) and MCF7 (**B**) cells transfected with siCELSR2 or the non-targeting control siNT. CREB-associated targets were selected from the MSigDB CREB_Q2 and CREB_Q4_01 gene sets. Row-wise z-scores were calculated across replicate samples and averaged within each condition. Red indicates higher abundance and blue/purple indicates lower abundance of the same corresponding protein. Proteins were categorized and hierarchically clustered. Mitochondrial metabolism includes proteins involved in respiratory-chain activity and assembly, mitochondrial translation, the TCA cycle, and amino-acid and fatty-acid metabolism. CELSR2 silencing was associated with a pronounced reduction in the relative abundance of mitochondrial metabolic proteins and CREB-associated targets in both cell lines.

**Supplementary Figure S9. CELSR2 promotes CREB activation and oxidative phosphorylation.** (**A**) Representative fluorescence images of HEK293T cells transfected with GFP or CELSR2-GFP expression plasmids. (**B**) Western blot analysis of GFP expression in HEK293T cells transfected with GFP, CELSR1-GFP, or CELSR2-GFP expression plasmids. Ponceau S staining served as the loading control. (**C**) Western blot analysis of CELSR2 expression in MDA-MB-468 cells transfected with siCtrl or *CELSR2*-targeting siRNAs. Ponceau S staining served as the loading control. (**D**) Western blot analysis of CELSR2, CREB, phospho-CREB (Ser133), mCherry, and β-tubulin expression in MGT19 cells stably expressing mCherry (mCh) or mCh-CELSR2. β-Tubulin served as the loading control. The P/no-P ratio corresponds to the abundance of phosphorylated CREB relative to total CREB. (**E**) Western blot analysis of CELSR2, ATP5A1, UQCRC2, SDHB, ATP5H, and β-tubulin expression in MGT19 cells stably expressing mCherry (mCh) or mCh+CELSR2. β-Tubulin served as the loading control.

**Supplementary Figure S10. *CELSR2* expression across normal breast tissues and breast cancer subtypes.** *CELSR2* mRNA expression in healthy breast tissues (GTEx), tumor-adjacent normal breast tissues (AT), and primary breast tumors from our gene expression database, including all breast cancers (all BC) and each molecular subtype (HR+/HER2−, HER2+, and TNBC). Expression values are shown as log₂-transformed normalized values. Box plots indicate the median and interquartile range. Statistical comparisons were performed between tumor-adjacent normal breast tissues (AT) and each breast cancer category (all BC, HR+/HER2−, HER2+, and TNBC) using two-sided t-tests. ns, not significant; ***P < 0.001; ****P < 0.0001.

**Table S1. List of siRNAs.**

**Table S2. List of plasmid constructs.**

**Table S3. List of shRNAs.**

**Table S4. Oligonucleotide primers used for RT-qPCR analyses.**

**Table S5. List of antibodies.**

**Table S6: List of gene expression data sets included in the study.**

**Table S7: Uni- and multivariate analyses for MFS in TNBC patients.**

---

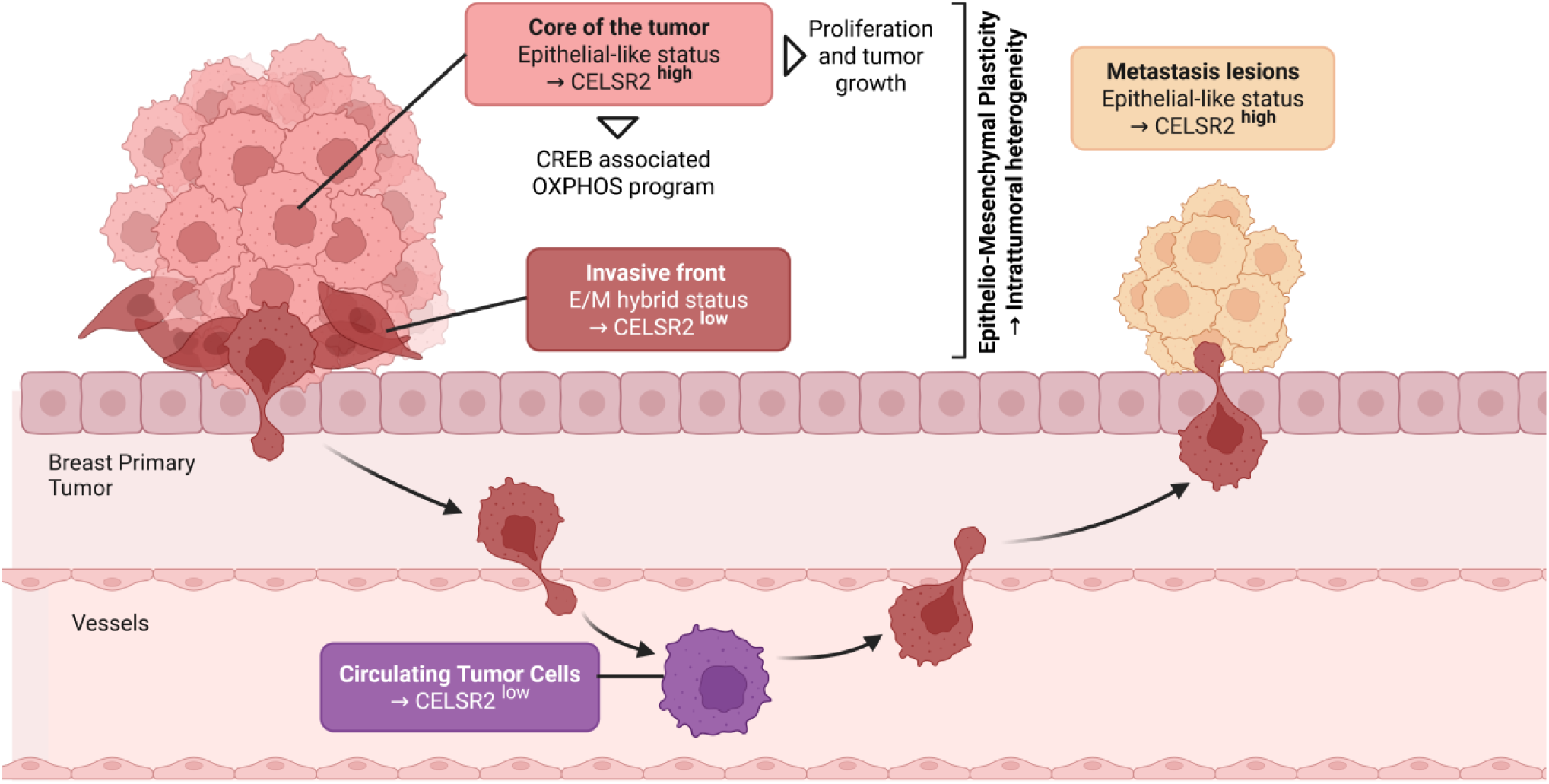

## Notes

### Competing Interest Statement

The authors have declared no competing interest.

