## Supplementary Figures S1-S10 for "The atypical cadherin CELSR2 regulates distinct epithelial–mesenchymal states and oxidative phosphorylation in triple-negative breast cancer"

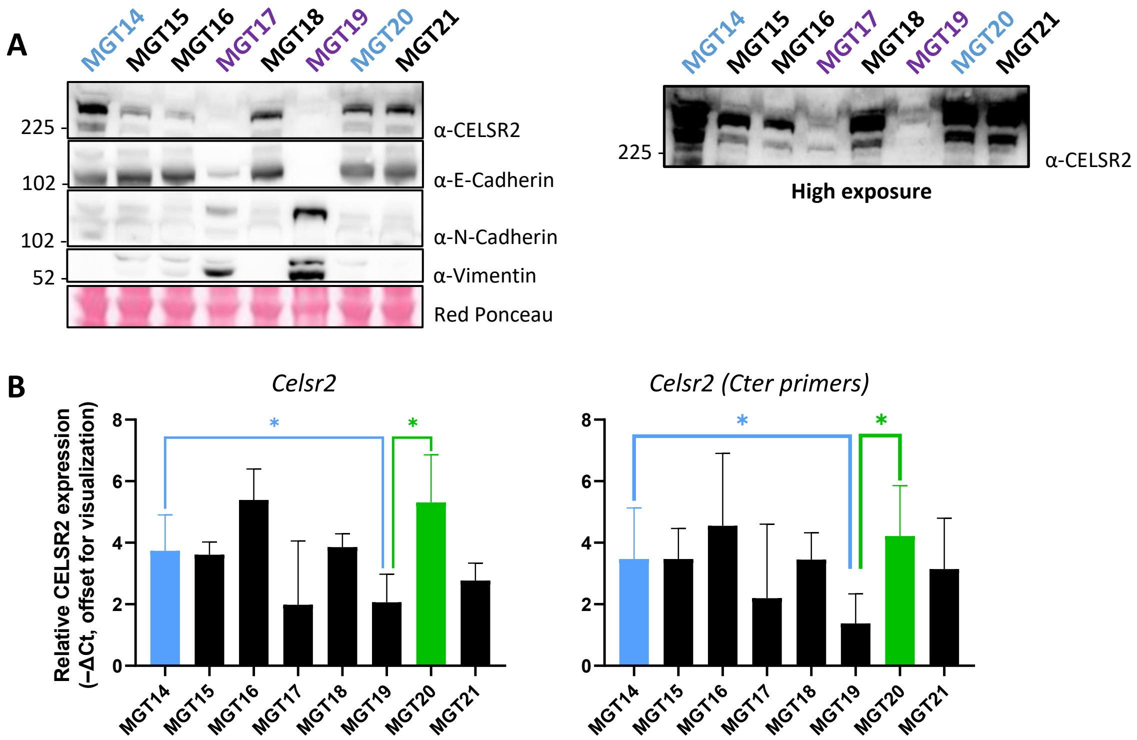


Koren et al, Supplementary Figure S1

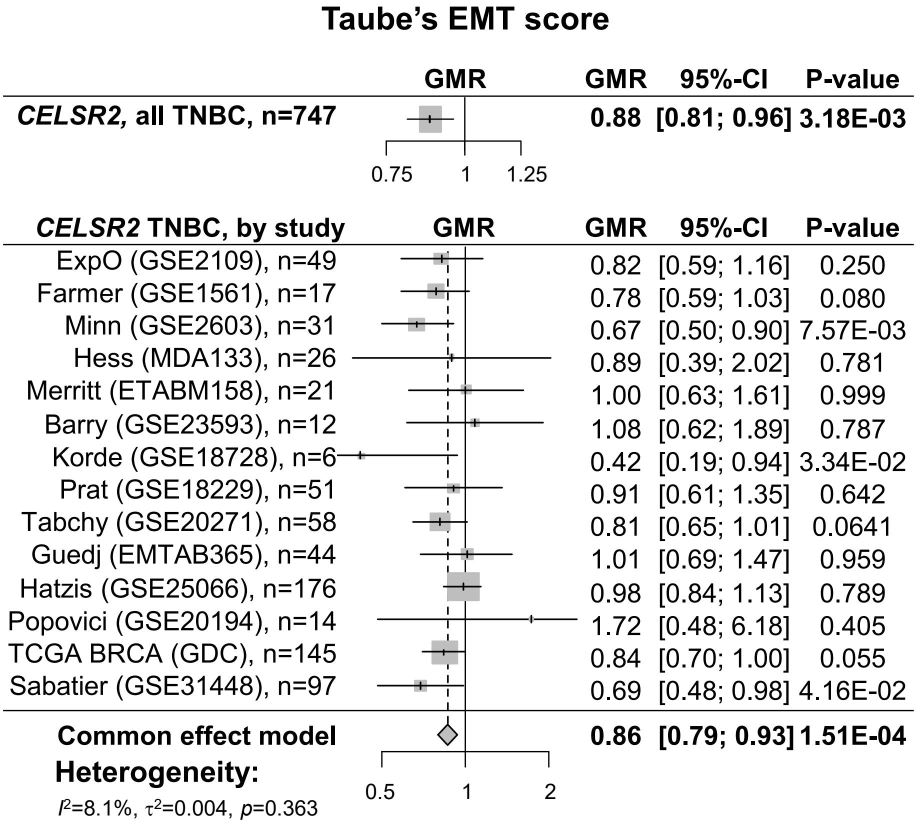


**Koren et al, Supplementary Figure S2**

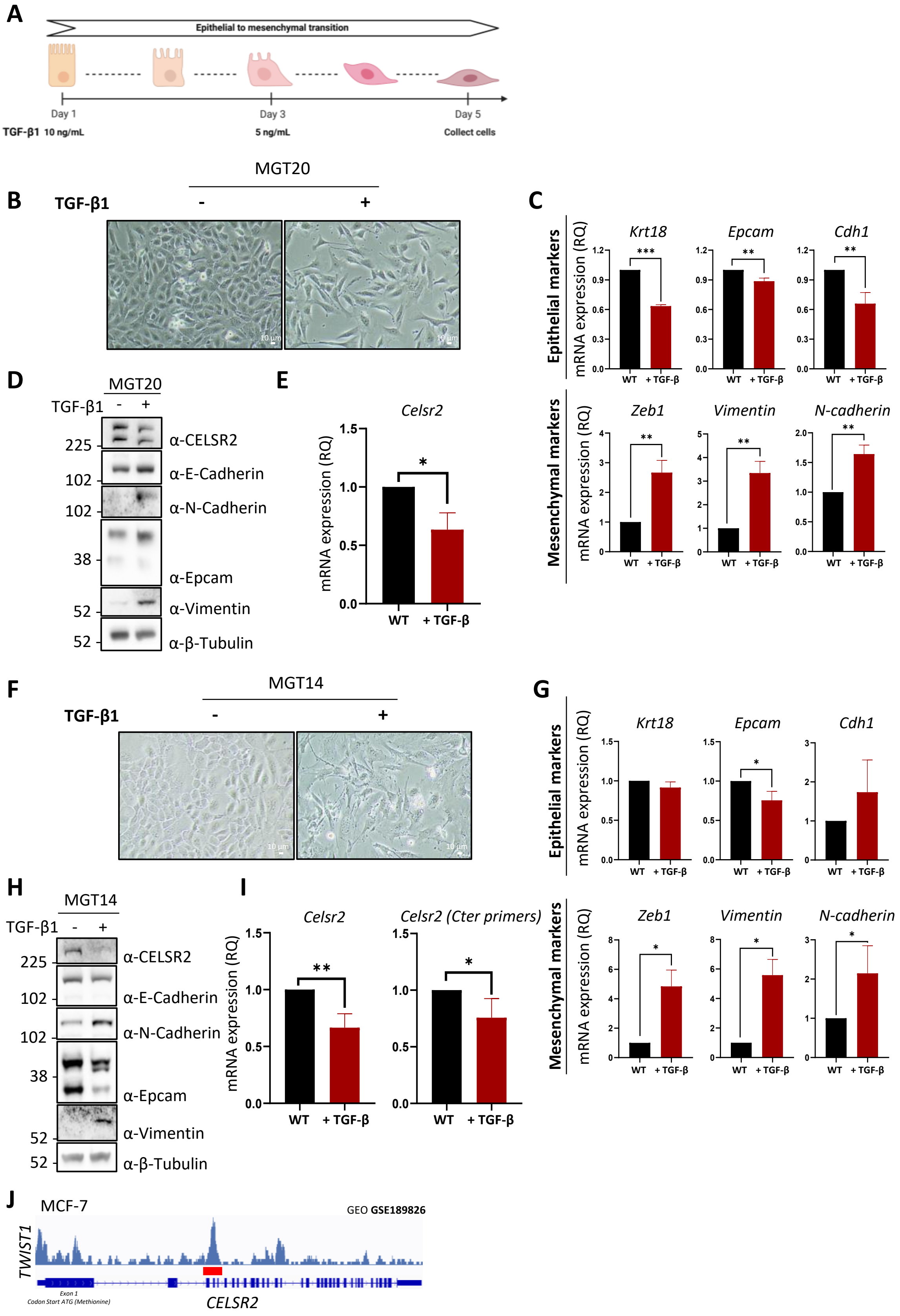


**Koren et al, Supplementary Figure S3**

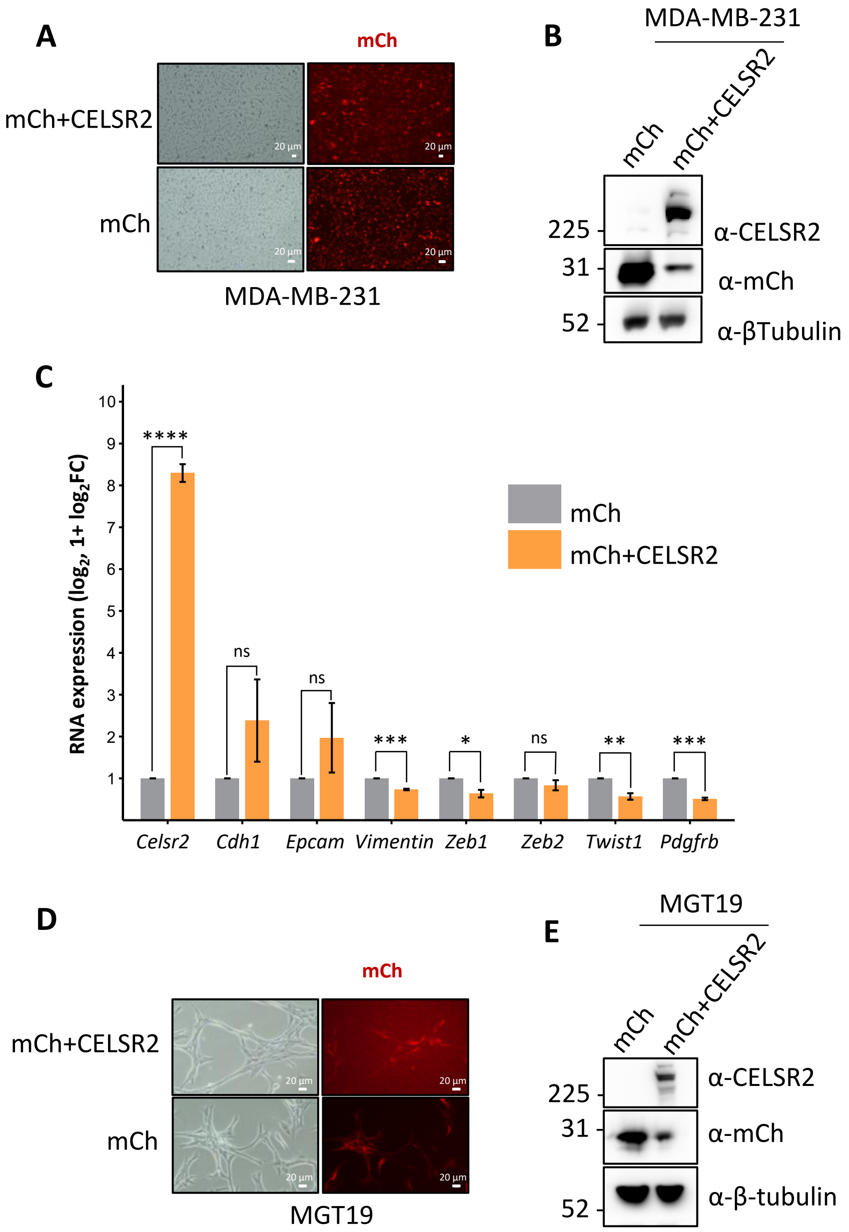


**Koren et al, Supplementary Figure S4**

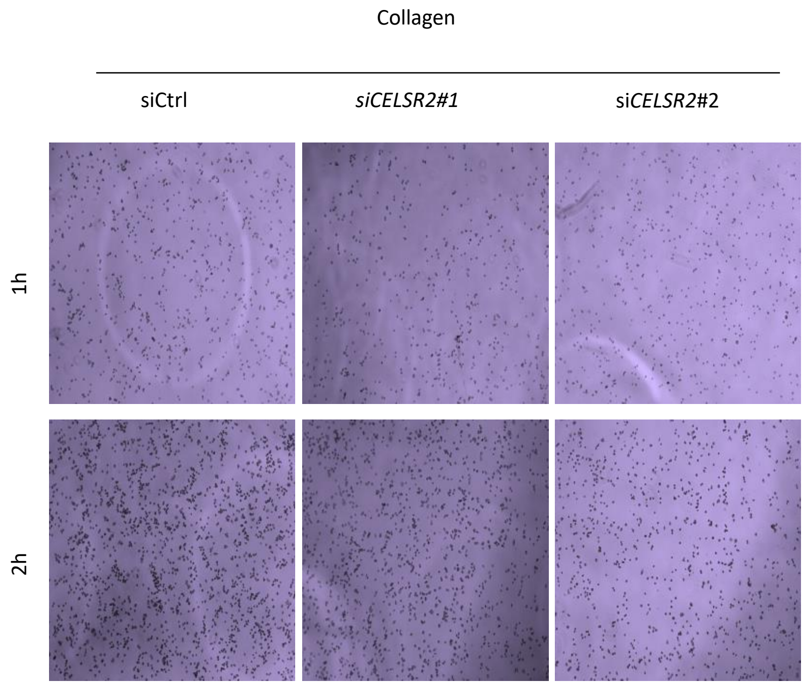


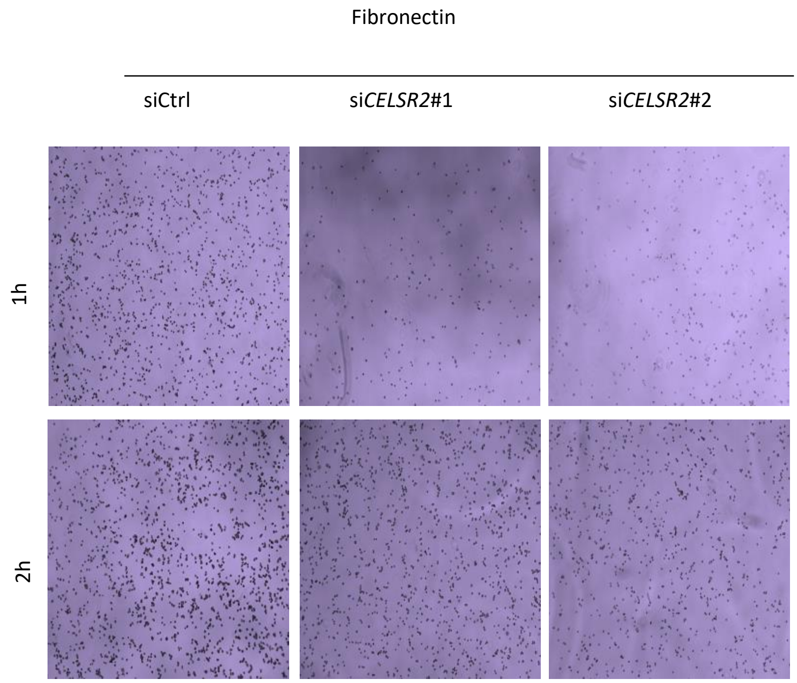


**Koren et al, Supplementary Figure S5**

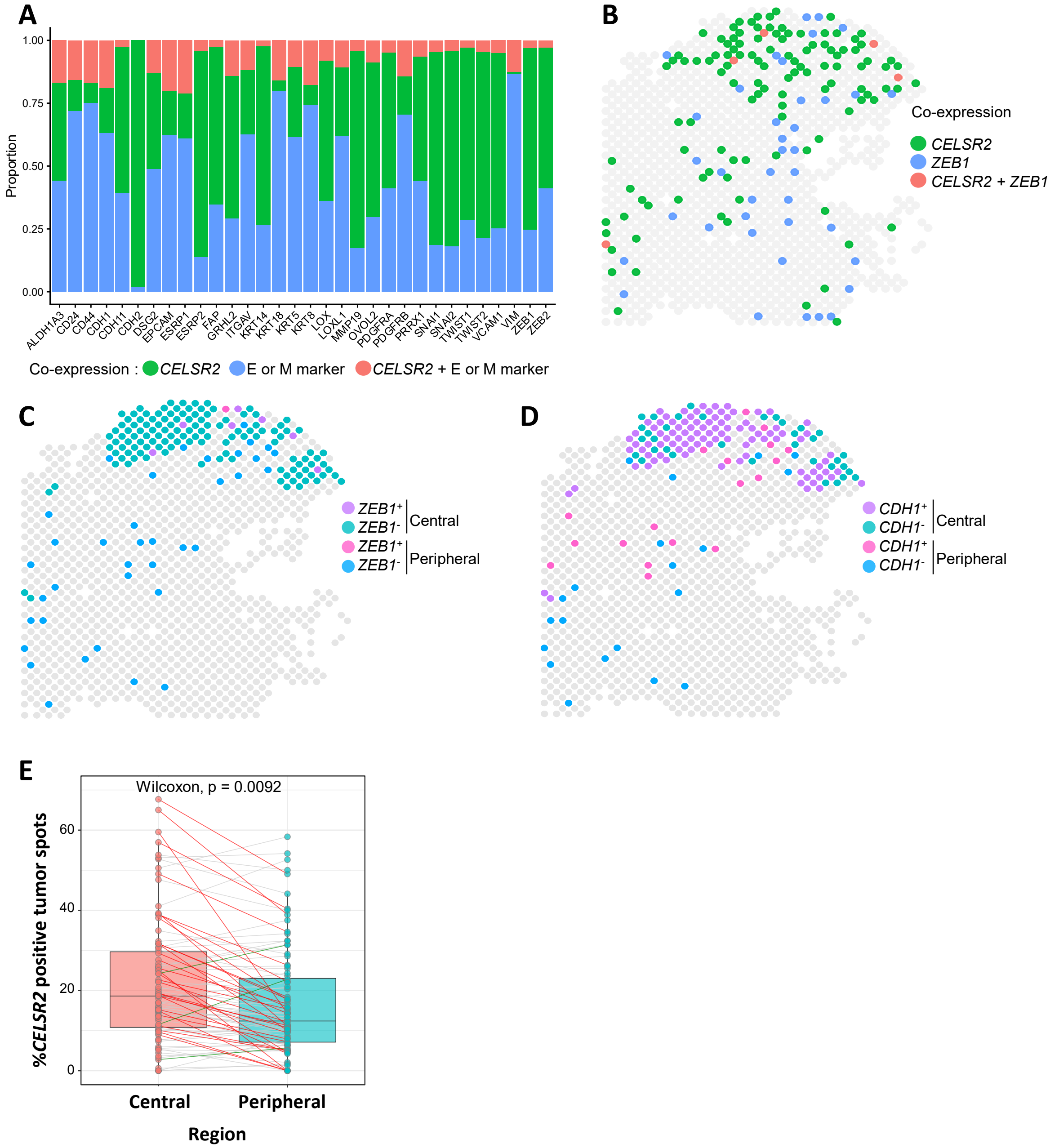


**Koren et al, Supplementary Figure S6**

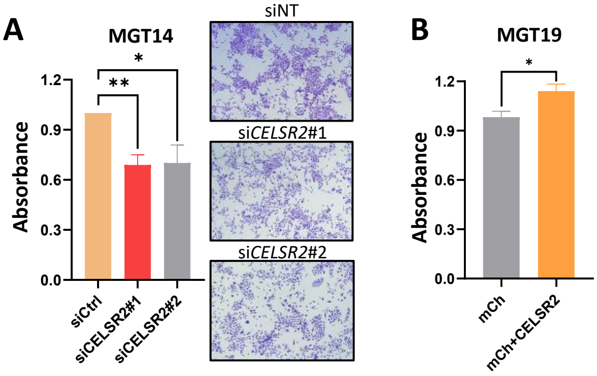


**Koren et al, Supplementary Figure S7**

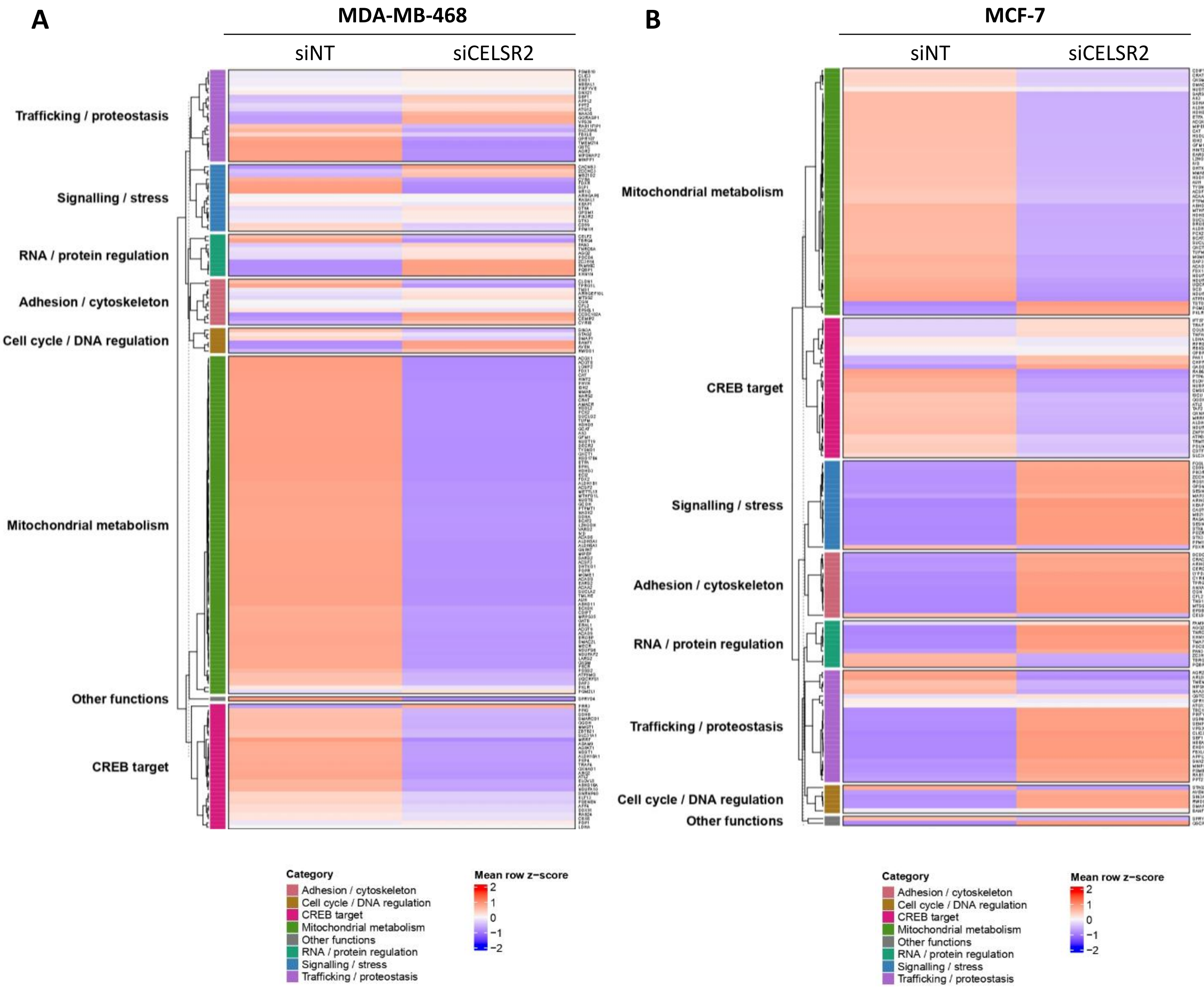


Koren et al, Supplementary Figure S8

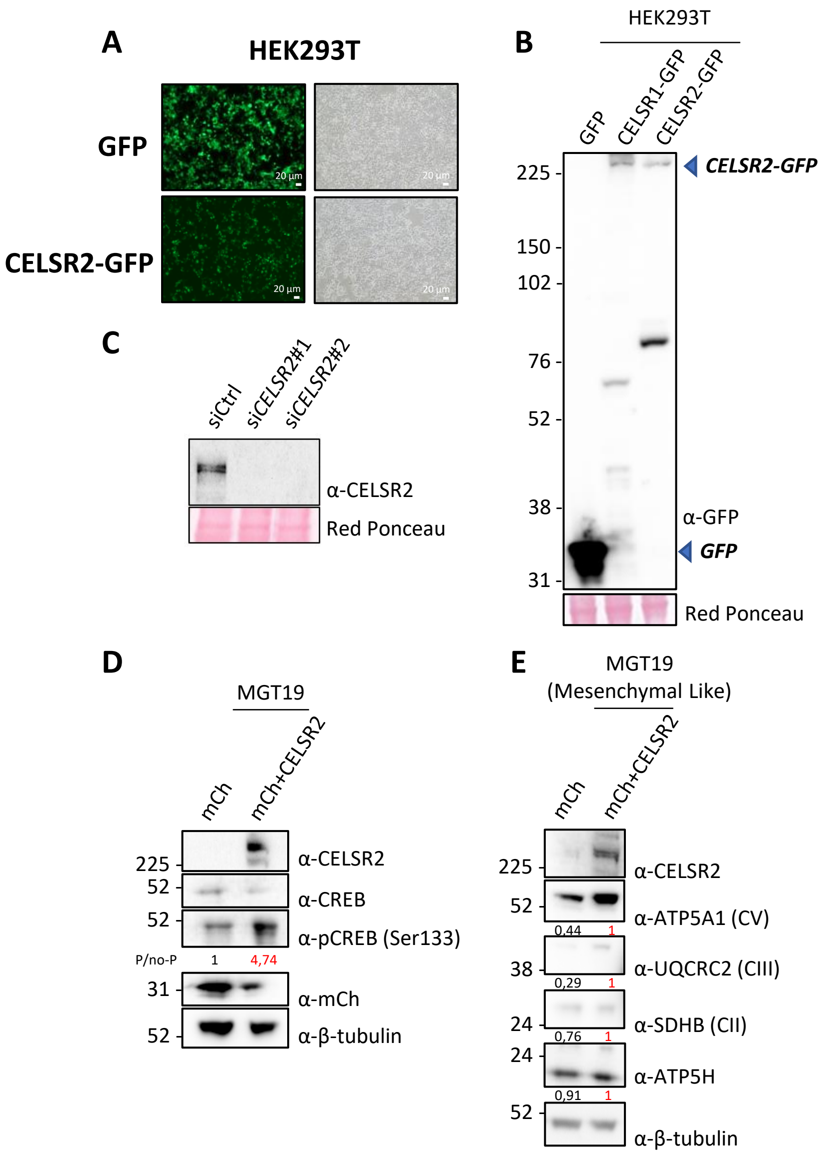


Koren et al, Supplementary Figure S9

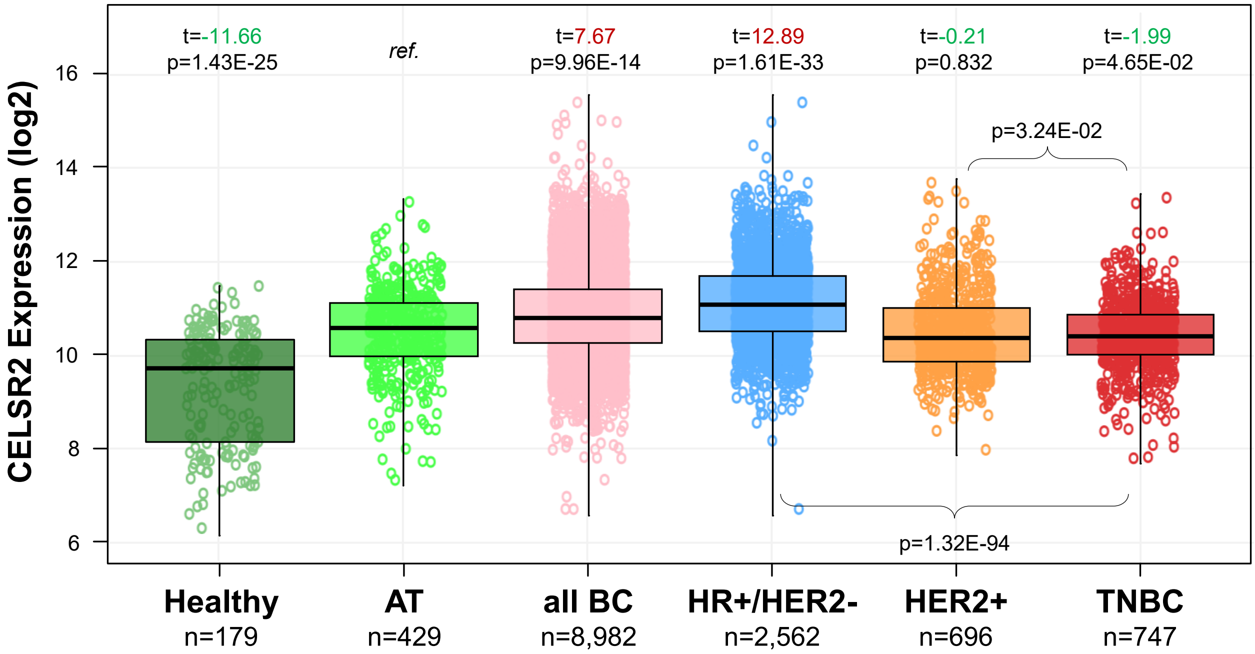


**Koren et al, Supplementary Figure S10**
