## Supplementary Tables S1-S7 for "The atypical cadherin CELSR2 regulates distinct epithelial–mesenchymal states and oxidative phosphorylation in triple-negative breast cancer"

**Table S1. siRNA.**

| **Target gene** | **Species** | **siRNA ID** | **Target sequence** | **Supplier** |
| --- | --- | --- | --- | --- |
| sihCELSR2#1 | Human | J-005460-06-0005 | GGGAGGAAGUUGAUUUCUA | Dharmacon |
| sihCELSR2#2 | Human | J-005460-07-0005 | GGUGAGCCCUCUUGACUAU | Dharmacon |
| simCELSR2#1 | Mouse | LQ-044500-06-0002 | GGACUGCACCUGAGCAAUA | Dharmacon |
| simCELSR2#2 | Mouse | LQ-044500-07-0002 | GUGAGAACGUGGCCCAGUA | Dharmacon |
| siNT |  | D-001810-01-50 | UGGUUUACAUGUCGACUAA | Dharmacon |

**.**

| **Plasmid** | **Source** |
| --- | --- |
| CELSR2-GFP | Gift from Tadashi Uemura |
| CELSR1-GFP | Gift from Danelle Devenport |
| PIV5.S157.mcherry | Addgene 36084 |
| PGK-CELSR2-T2A-mCherry | Azenta Life Science |
| GW-pRLL-luc2-GFP | TrGET Platform, CRCM |
| CRE luciferase reporter | Gift from Ines Liebscher (Promega) |
| Renilla luciferase reporter | Gift from Ines Liebscher (Promega) |
| psPAX2 packaging plasmid | Addgene |
| pMD2.G (VSV-G envelope plasmid) | Addgene |

**Table S2. Plasmids.**

**Table S3. ShRNA.**

| **shRNA ID** | **Species** | **Target sequence** | **Supplier** |
| --- | --- | --- | --- |
| shCELSR2#1 | Human | TAGGATTCCACCTCTTCCC | Horizon Discovery |
| shCELSR2#2 | Human | TTTACATTCCTTTTCCGAC | Horizon Discovery |
| shCtrl |  |  | Horizon Discovery |

| **Gene** | **Species** | **Forward primer** | **Reverse primer** |
| --- | --- | --- | --- |
| *CELSR2 (NTF)* | Human | 5’GCTACATCCCCTTCTTGCTG3’ | 5’TCCTCTTCCTCCTCCTCTTC3’ |
| *VIM* | Human | 5'AGGCAAAGCAGGAGTCCACTGA3' | 5'ATCTGGCGTTCCAGGGACTCAT3' |
| *KRT19* | Human | 5'AGCTAGAGGTGAAGATCCGCGA3' | 5'GCAGGACAATCCTGGAGTTCTC3' |
| *EPCAM* | Human | 5'GCTGGCCGTAAACTGCTTTGT3' | 5'TGCCTTCATCACCAAACATTTGGC3' |
| *CDH1* | Human | 5’CCCACCACGTACAAGGGTC3’ | 5’CTGGGGTATTGGGGGCATC3’ |
| *CDH2* | Human | 5’CCTCCAGAGTTTACTGCCATGAC3’ | 5’GTAGGATCTCCGCCACTGATTC3’ |
| *PDGFRA* | Human | 5'GTGATAATCCCCACAGGCACA3' | 5'ACATGAACAGGGGCATTCGT3' |
| *GAPDH* | Human | 5’GTCTCCTCTGACTTCAACAGCG3’ | 5’ACCACCCTGTTGCTGTAGCCAA3’ |
| *KI67* | Human | 5′GAGGTGTGCAGAAAATCCAAA3′ | 5′CTGTCCCTATGACTTCTGGTTGT3′ |
| *VCAM1* | Human | 5’ TTTGACAGGCTGGAGATAGACT3’ | 5’TCAATGTGTAATTTAGCTCGGCA3’ |
| *Celsr2 (NTF)* | Mouse | 5’TGCACGTCAGTCACCTTCTC3’ | 5’CTGGTAGGCCACCTTGACAT3’ |
| *Celsr2 (Cter primers)* | Mouse | 5’ATAGTGGTAGTGGGCCCCTT3’ | 5’GACTGCCCTCACTGATGGAG3’ |
| *Krt18* | Mouse | 5’GTGCCACCATTGACAACTCC3’ | 5’AATCCACCTCCACACTGACC3’ |
| *Cdh1* | Mouse | 5’CAGGTCTCCTCATGGCTTTGC3’ | 5’CTTCCGAAAAGAAGGCTGTCC3’ |
| *Vim* | Mouse | 5’GATGGCTCGTCACCTTCGTG3’ | 5’GAAATCCTGCTCTCCTCGCC3’ |
| *Zeb1* | Mouse | 5’GCTGGCAAGACAACGTGAAAG3’ | 5’GCCTCAGGATAAATGACGGC3’ |
| *Zeb2* | Mouse | 5’GGGATCACTCGATGGACGAC3’ | 5’GCCCCTTCTGTCCCTCTCTA3’ |
| *Epcam* | Mouse | 5’ACCTGAGAGTGAACGGAGAGCC3’ | 5’TGCATGGAGAACTCGGGTGCCT3’ |
| *Cdh2* | Mouse | 5’AGCGCAGTCTTACCGAAGG3’ | 5’TCGCTGCTTTCATACTGAACTTT3’ |
| *Twist1* | Mouse | 5’CCGGAGACCTAGATGTCATTGT3’ | 5’CCACGCCCTGATTCTTGTGA3’ |
| *Pdgfrb* | Mouse | 5’CTTGTTCTGGGACGCACTCT3’ | 5’TTGTTCCGGTGCAGGTAGTC3’ |

**Table S4. Oligonucleotide primers for RT-qPCR analyses**

**Table S5. Antibodies for Western Blot and immunofluorescence.**

| **Target** | **Host** | **Clonality** | **Application** | **Dilution** | **Supplier** | **Catalog#** | **RRID** |
| --- | --- | --- | --- | --- | --- | --- | --- |
| CELSR2 | Rabbit | Monoclonal | WB | 1 : 1000 | Cell Signaling | 47061 | AB_2799319 |
| NDUFV2 | Rabbit | Polyclonal | WB | 1 : 5000 | Proteintech | 15301-1-AP | AB_2149048 |
| NUBPL | Rabbit | Polyclonal | WB | 1 : 1000 | Proteintech | 17393-1-AP | AB_2878402 |
| ATP5H | Rabbit | Polyclonal | WB | 1 : 2000 | Proteintech | 17589-1-AP | AB_2062046 |
| CREB | Rabbit | Monoclonal | WB | 1 : 1000 | Cell Signaling | 9197 | AB_331277 |
| pCREB (Ser133) | Rabbit | Monoclonal | WB | 1 : 1000 | Cell Signaling | 9198 | AB_2561044 |
| OXPHOS complex (I–V cocktail) | Mouse | Monoclonal | WB | 1 : 1000 | abcam | ab110413 | AB_2629281 |
| E-cadherin | Rabbit | Monoclonal | WB | 1 : 1000 | Cell Signaling | 3195 | AB_2291471 |
| N-cadherin | Rabbit | Monoclonal | WB | 1 : 1000 | Cell Signaling | 13116 | AB_2687616 |
| Vimentin | Rabbit | Monoclonal | WB | 1 : 1000 | Cell Signaling | 5741 | AB_10695459 |
| β-tubulin | Mouse | Monoclonal | WB | 1 : 1000 | Cell Signaling | 86298 | AB_2715541 |
| Epcam | Rabbit | Monoclonal | WB | 1 : 1000 | Cell Signaling | 42515 | AB_3739901 |
| mCh | Rabbit | Polyclonal | WB | 1 : 1000 |  |  |  |
| GFP | Rabbit | Polyclonal | WB | 1 : 2000 | Thermo Fisher Scientific | A-11122 | AB_221569 |
| KRT18 | Mouse | Monoclonal | WB | 1 : 1000 | Santa Cruz Biotechnology | sc-6259 | AB_627850 |
| ITGA6 | Rabbit | Polyclonal | WB | 1 : 1000 | Bethyl Laboratories | A303-682A | AB_11204419 |
| ITGB4 | Rabbit | Polyclonal | WB | 1 : 1000 | Thermo Fisher Scientific | PA5-83325 | AB_2790481 |
| ITGAV | Mouse | Monoclonal | WB | 1 : 1000 | BD Biosciences | 611012 | AB_2129503 |
| ITGB1 | Rabbit | Polyclonal | WB | 1 : 1000 | Bethyl Laboratories | A303-735A | AB_11204764 |
| E-cadherin | Goat | Polyclonal | IF | 1 : 500 | R and D Systems | AF648 | AB_355504 |
| ZO-1 | Mouse | Monoclonal | IF | 1 : 200 | Thermo Fisher Scientific | 33-9100 | AB_87181 |
| CELSR2 | Rabbit | Polyclonal | IF | 1 : 500 | Cloud Clone Corporation | PAG433Mu01 |  |
| Donkey anti-  Rabbit IgG  (H+L) 488 | Donkey | Polyclonal | IF | 1 : 2000 | Alexa FluorTM,  ThermoFisher | R37118 | AB_2556546 |
| Donkey anti-  Goat IgG  (H+L) 647 | Donkey | Polyclonal | IF | 1 : 2000 | Alexa FluorTM,  ThermoFisher | A32849TR | AB_2866498 |
| Donkey anti-  Mouse IgG  (H+L) 546 | Donkey | Polyclonal | IF | 1 : 2000 | Alexa FluorTM,  ThermoFisher | A10036 | AB_11180613 |

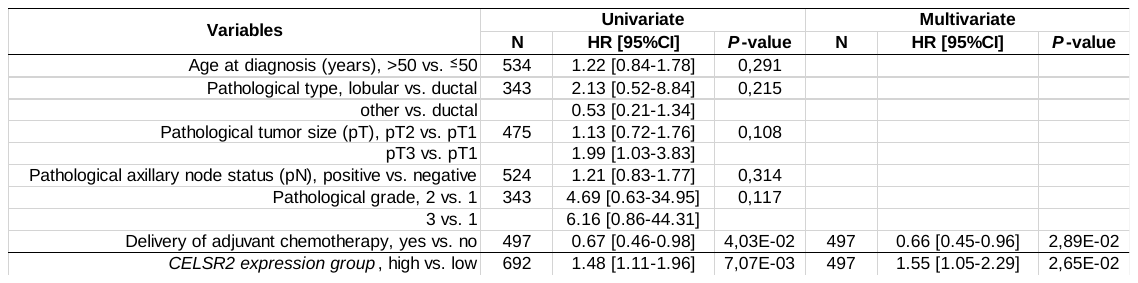
**Table S6. List of gene expression data sets included in the study.**

| **Reference** | **Source of data** | **Technological platform** | | **N° of primary cancer samples** | **N° of normal breast samples from** | |
| --- | --- | --- | --- | --- | --- | --- |
|  |  |  |  |  | **Cancer patients** | **Healthy**  **people** |
| van de Vijver et al.,  NEJM 2002 | http://microarray-pubs.stanford.edu/wound_NKI/ | Agilent Hu25K | | 254 |  |  |
| van't Veer et al.,  Nature 2002 | http://www.rii.com/publications/2002/vantveer.html | Agilent Hu25K | | 117 |  |  |
| Expression Project for Oncology (expO), 2005 | https://expo.intgen.org/geo GEO: GSE2109 | Affymetrix U133 Plus 2.0 | | 348 |  |  |
| Farmer P et al.,  Oncogene 2005 | GEO: GSE1561 | Affymetrix U133A | | 49 |  |  |
| Minn AJ et al.,  Nature 2005 | GEO: GSE2603 | Affymetrix U133A | | 99 |  |  |
| Wang Y et al.,  Lancet 2005 | GEO: GSE2034 | Affymetrix U133A | | 286 |  |  |
| Hess KR et al.,  J Clin Oncol 2006 | MDA133 | Affymetrix U133A | | 131 |  |  |
| Ivshina et al.,  Cancer Res 2006 | GEO: GSE4922, GSE1456 | Affymetrix U133 A+B | | 448 |  |  |
| Sotiriou C et al.,  J Natl Cancer Inst 2006 | GEO: GSE2990 | Affymetrix U133A | | 80 |  |  |
| Bonnefoi et al.,  Lancet Oncol 2007 | GEO: GSE6861, GSE4779 | Affymetrix X3P | | 125 |  |  |
| Desmedt C etal.,  Clin Cancer Res 2007 | GEO: GSE7390 | Affymetrix U133A | | 154 |  |  |
| Miller WR et al.,  Breast Cancer Res 2010 | GEO: GSE5462 | Affymetrix U133A | | 116 |  |  |
| Klein A et al.,  Int J Cancer 2007 | GEO: GSE6596 | Affymetrix U133A | | 24 | 2 |  |
| Bos et al., Nature 2009 | GEO: GSE12276 | Affymetrix U133 Plus 2.0 | | 204 |  |  |
| Hoeflich et al.,  Clin Cancer Res 2009 | GEO: GSE12763 | Affymetrix U133 Plus 2.0 | | 30 |  |  |
| Marty et al.,  Breast Cancer Res 2008 | GEO: GSE13787 | Affymetrix U133 Plus 2.0 | | 23 |  |  |
| Merritt WM et al.,  N Engl J Med 2008 | Array Express: E-MTAB-158 | Affymetrix U133AAofAv2 | | 130 |  |  |
| Schmidt M etal.,  Cancer Res 2008 | GEO: GSE11121 | Affymetrix U133A | | 200 |  |  |
| Yu K et al.,  PLoS Genet 2008 | GEO: GSE5364 | Affymetrix U133A | | 183 | 13 |  |
| Zhang Y et al.,  Breast Cancer Res Treat 2009 | GEO: GSE12093 | Affymetrix U133A | | 136 |  |  |
| Barry et al.,  J Clin Oncol 2010 | GEO: GSE23593 | Affymetrix U133 Plus 2.0 | | 50 |  |  |
| Iwamoto T et al.,  J Natl Cancer Inst 2011 | GEO: GSE22093, GSE22597 | Affymetrix U133A | | 100 |  |  |
| Korde et al.,  Breast Cancer Res Treat 2010 | GEO: GSE18728 | Affymetrix U133 Plus 2.0 | | 61 |  |  |
| Prat A et al.,  Breast Cancer Res 2010 | GEO: GSE18229 | Agilent Hu25K | | 264 | 14 |  |
| Silver et al.,  J Clin Oncol 2010 | GEO: GSE18864 | Affymetrix U133 Plus 2.0 | | 84 |  |  |
| Tabchy A et al.,  Clin Cancer Res 2010 | GEO: GSE20271 | Affymetrix U133A | | 178 |  |  |
| Jonsson et al.,  BCR 2010 | GEO: GSE22133 | Swegene H_v2.1.1 55K | | 346 |  |  |
| Chen et al.,  Breast Cancer Res Treat 2010 | GEO: GSE10780 | Affymetrix U133 Plus 2.0 | | 42 | 143 |  |
| Desmedt et al.,  J Clin Oncol 2011 | GEO: GSE16446 | Affymetrix U133 Plus 2.0 | | 120 |  |  |
| Guedj et al.,  Oncogene 2011 | Array Express: E-MTAB-365 | Affymetrix U133 Plus 2.0 | | 452 |  |  |
| Hatzis C et al.,  JAMA 2011 | GEO: GSE25066 | Affymetrix U133A | | 504 |  |  |
| Popovici V et al.,  Breast Cancer Res 2010 | GEO: GSE20194 | Affymetrix U133A | | 91 |  |  |
| TCGA,  Nature 2012 | TCGA Data Portal - BRCA - | Illumina, RNAseq V2 | | 1,092 | 113 |  |
| Ellis et al.,  Nature 2012 | GEO: GSE29442, GSE35186 | Agilent-014850 4x44K | | 201 | 144 |  |
| Curtis et al.,  Nature 2012 | EGA: EGAS00000000083 | Illumina HT 12 | | 1,974 |  |  |
| Sabatier R et al., (our IPC series) PLoS One 2011 | GEO: GSE31448 | Affymetrix U133 Plus 2.0 | | 286 |  |  |
| The GTEx Consortium, Nat Genet. 2013 | [https://xena.ucsc.edu/public/](https://www.gtexportal.org/) | Illumina, RNAseq V2 | |  |  | 179 |
| TOTAL | | |  | 8,982 | 429 | 179 |

**Table S7. Uni- and multivariate analyses for MFS in TNBC patients.**

| **Variables** | **Univariate** | | | **Multivariate** | | |
| --- | --- | --- | --- | --- | --- | --- |
|  | **N** | **HR [95%CI]** | ***P*-value** | **N** | **HR [95%CI]** | ***P*-value** |
| Age at diagnosis (years), >50 vs. ≤50 | 534 | 1.22 [0.84-1.78] | 0.291 |  |  |  |
| Pathological type, lobular vs. ductal | 343 | 2.13 [0.52-8.84] | 0.215 |  |  |  |
| other vs. ductal |  | 0.53 [0.21-1.34] |  |  |  |  |
| Pathological tumor size (pT), pT2 vs. pT1 | 475 | 1.13 [0.72-1.76] | 0.108 |  |  |  |
| Pathological tumor pT3 vs. pT1 |  | 1.99 [1.03-3.83] |  |  |  |  |
| Pathological axillary node status (pN), positive vs. negative | 524 | 1.21 [0.83-1.77] | 0.314 |  |  |  |
| Pathological grade, 2 vs. 1 | 343 | 4.69 [0.63-34.95] | 0.117 |  |  |  |
| 3 vs. 1 |  | 6.16 [0.86-44.31] |  |  |  |  |
| Delivery of adjuvant chemotherapy, yes vs. no | 497 | 0.67 [0.46-0.98] | 4.03E-02 | 497 | 0.66 [0.45-0.96] | 2.89E-02 |
| *CELSR2 expression group*, high vs. low | 692 | 1.48 [1.11-1.96] | 7.07E-03 | 497 | 1.55 [1.05-2.29] | 2.65E-02 |
